# The regulatory role of anti-sigma factor, RsbW, in *Clostridioides difficile* stress response, persistence and infection

**DOI:** 10.1101/2022.08.22.504775

**Authors:** Jeffrey K. J. Cheng, Tanja Đapa, Ivan Y. L. Chan, Thomas MacCreath, Ross Slater, Meera Unnikrishnan

## Abstract

**Summary:** The anaerobic pathogen *Clostridioides difficile*, a primary cause of antibiotic-associated diarrhoea, faces a variety of stresses in the environment and in the mammalian gut. To cope with environmental stresses, it uses the alternative sigma factor B (σ^B^) to modulate gene transcription, which is regulated by an anti-sigma factor, RsbW. To understand the role of RsbW in *C. difficile* physiology, a *rsbW* mutant (Δ*rsbW*) where σ^B^ is ‘always on’, was generated. Δ*rsbW* did not have deleterious fitness defects but tolerated acidic environments and detoxified reactive oxygen and nitrogen species better. Δ*rsbW* was defective in spore and biofilm formation, adhered better to human gut epithelia and was less virulent in a *Galleria mellonella* infection model. A transcriptomic analysis to understand this unique phenotype showed a change in expression of some σ^B^-controlled genes along with several non-σ^B^ controlled genes. Interestingly, sinRR’ locus that encodes a pleiotropic regulator, was highly upregulated in Δ*rsbW* indicating a potential indirect role for σ^B^ or RsbW in control of *sinRR’*. Furthermore, the unexpected lower intracellular levels of σ^B^ observed suggest post translational control mechanisms. Our study thus provides new insight into the regulatory role of RsbW and the complexity of regulatory networks in *C. difficile*.

**Importance:** Pathogens, like *C. difficile*, face a range of stresses in the environment and within the host. Alternate transcriptional factors such as sigma factor B (σ^B^) enables the bacterium to respond quickly to different types of stresses and are conserved across bacteria. Anti-sigma factors like RsbW control the activation of genes via these pathways. Such transcriptional control systems provide pathogens like *C. difficile* a route to tolerance and detoxification of harmful compounds. In this study we investigate the role of RsbW in *C. difficile* physiology. We demonstrate distinctive phenotypes for a *rsbW* mutant in growth, persistence and virulence. Our data suggest new σ^B^ regulatory circuits in *C. difficile*. Understanding bacterial responses to external stress is key to designing better strategies to combat this highly resilient bacterial pathogen.

## Introduction

*Clostridioides difficile* is a Gram-positive, spore-forming, anaerobic bacterium found ubiquitously in nature in soil, water and nosocomial environments. Transmission primarily occurs through the ingestion of spores via the faecal-oral route in healthcare and community settings. *C. difficile* has the opportunity to proliferate and induce symptoms of varying severity when the normal microbiota in the gastrointestinal (GI) tract is disrupted, usually as a result of antibiotic treatment. *C. difficile* infection (CDI) is typically associated with severe diarrhoea, pseudomembranous colitis and toxin megacolon^1,2^. The greater awareness for CDI, has reduced cases of healthcare-associated infections and in-hospital deaths, and has shifted more attention to recurrent infections^3^. High rates of recurrent CDI at rates of up to 30%^4,5^ have increased the cost burden on healthcare systems worldwide. One of the reasons for high rates of recurrence is the ability of *C. difficile* to persist in extreme environmental conditions, both in hospital and community environments and within the host gut.

*C. difficile* spores are known to resist a range of environmental stresses including human stomach acid^6,7^, however upon germination, the vegetative cells are exposed to a myriad host-derived stressors. Each of the GI tract compartments like the duodenum, jejunum, ileum present *C. difficile* with varying levels of oxygen concentrations, pH, osmolarity, immune responses and inter-species competition^8,9^. Like many other Firmicutes, *C. difficile* possesses alternative sigma factors to enable it to respond against such external stimuli, one example being sigma factor B (σ^B^)^10^. By changing the transcriptional target of the RNA polymerase, the bacterium can express a plethora of general stress response genes to ensure its adaptation and survival.

σ^B^ has been best characterised in *Bacillus subtilis*, where 7 other different proteins are involved in the activation of σ^B^ through the partner-switching complex, phosphatases and a transducer stressosome complex^11–15^. The number of alternative σ factors and ‘Rsb’ proteins appear to differ between species, suggesting that this diversity can be associated with the variety of stresses encountered in its lifecycle^16^. In *C. difficile*, only the partner-switching complex and a PP2C-type phosphatase has been uncovered^17,18^. The genes encoding the post-translational partner switching modules (RsbV and RsbW) are always in the σ^B^ operon and, currently, appear to be universally conserved in all strains expressing σ^B^ (Figure S1A). In this post-translational paradigm, σ^B^ can exist in two states, unbound and bound. During normal growth in the absence of stress, anti-sigma factor RsbW sequesters σ^B^ to prevent unnecessary transcription^11,19,20^. Only through the dephosphorylation of anti-anti-sigma factor RsbV, can RsbW preferentially release σ^B^ in favour for RsbV^21^. This event is mediated by a serine/threonine protein phosphatase 2C, RsbZ, and subsequently controls σ^B^ activation^18^. The balance is restored through the additional activity of RsbW, acting as a kinase, it phosphorylates RsbV and returns to restrain σ^B 22^ (Figure S1B).

The roles of σ^B^ and associated Rsb proteins has been studied recently in *C. difficile*^17,18,23,24^, and RsbW was shown to directly bind to σ^B^. From RNA-seq studies, σ^B^ was involved primarily in oxidative and nitrosative stress response, detoxification of reactive oxidative species (ROS) and reactive nitrogen species (RNS) was successfully demonstrated in subsequent phenotypic assays^17,18,23^. σ^B^ has also been implicated in low oxygen tolerance, protection from low pH, DNA repair, tellurite and thiol resistance^17,18^. Furthermore, σ^B^ mutants displayed increased sporulation frequency and higher colonisation in a murine infection model^17^.

Unlike σ^B^, the role of the anti-sigma factor RsbW during bacterial growth or under conditions of stress has been understudied. One of the reasons for this is because in many species, σ^B^ is autoregulated, causing uncontrolled transcription and a deleterious phenotype in the absence of RsbW, as shown in *Bacillus*^11^. In *C. difficile* as it was presumed that a deletion mutant would be toxic^20,25^, only an artificial overexpressing strain has been examined^18^. However, the *C. difficile* σ^B^ operon appears to be controlled only by housekeeping σ^A^, unlike in other species, as a σ^B^ promoter has not been identified immediately upstream of the operon^26–28^. This indicates that the transcription of the *sigB* would be unaltered in the absence of RsbW.

In this study we created a *rsbW* deletion mutant in *C. difficile* R20291, to explore the regulatory roles of RsbW. In this strain, the lack of a partner-switching mechanism would result in unbound σ^B^, and hence a ‘constitutive’ expression of σ^B^. Here we demonstrate the lack of a fitness defect in the Δ*rsbW* mutant but better survival in different stress-inducing conditions. The absence of RsbW also impacted cell adhesion and virulence. Interestingly, a RNAseq analysis revealed transcriptional changes distinct to those previously reported for a σ^B^ mutant and an upregulation of the pleiotropic regulator SinRR’. Examination of relative intracellular concentrations of σ^B^ revealed a decreased level of σ^B^ indicating posttranslational control. The results from this study demonstrate the role of RsbW in modulating stress responses and virulence and indicate new mechanisms of σ^B^ control.

## Experimental Procedures

### Bacterial Strains and Culture Conditions

The bacterial strains and plasmids used in this study are listed in the Table S1. *C. difficile* strains were grown in pre-reduced Brain-Heart Infusion (BHI) with 0.5% (w/v) yeast-exact and 0.1% (w/v) L-cysteine (BHI-S) or Tryptose Yeast (TY). Media were also supplemented antimicrobials; cefoxitin (8 μg/mL), D-cycloserine (250 μg/mL), thiamphenicol (15 μg/mL) where required. Complemented strains were induced with anhydrotetracycline (20 ng/mL) for the *Ptet* promoter containing pRPF185-derivatives in this study^68^. Cultures were grown under anaerobic conditions (80% N_2_, 10% CO_2_ and 10% H_2_) in a Don Whitely Scientific MG500 Anaerobic Workstation (Yorkshire, UK). *Escherichia coli* strains were grown in LB media supplemented with chloramphenicol (15 μg/mL) where necessary.

### Construction of Δ*rsbW*

The allele exchange system from Cartman *et al*., was employed in the generation of the Δ*rsbW* mutant in *C. difficile* strain R20291. The specific details of the generation of mutant and complement strain can be found in the Supplementary Methods. The allelic exchange cassette was generated “in frame”, flanking the gene of interest and cloned into the pMTL-SC7315 vector. The construct was transformed into *E. coli* DH5α by heat shock, then electrocompetent *E. coli* donor strain CA434 and subsequently conjugated into R20291. Transconjugants, first and second cross-over mutants were selected with agar plates with/without the appropriate antibiotics and supplements. Sanger sequencing confirmed successful generation of mutants.

### Fitness and Overexpression Assay

Overnight (O/N) cultures of bacteria were diluted to an OD_600_ of 0.05 in a large volume pre-reduced BHI-S and TY broth, grown in anaerobic conditions and OD_600_ was measured on a Novaspec Pro spectrophotometer (Biochrom, US). Overexpression of rsbW was induced with 200, 1000 and 2000 ng/mL of anhydrotetracycline (aTc) in BHI-S broth supplemented with thiamphenicol. The viability of bacteria and plasmid stability was also measured via a spot assay on BHI-S plates supplemented with/without thiamphenicol and/or aTc; O/N cultures were standardised to an OD_600_ of 1.0, serially diluted in pre-reduced sterile phosphate buffered saline (PBS) and 10 μL of each dilution were dispensed onto agar plates (150 × 150 × 15 mm, Sarstedt, Germany). Colony forming units were counted after incubation for 24 h in anaerobic conditions.

### Oxygen Tolerance Assay

To measure tolerance and growth in an aerobic environment, strains were grown in soft agar tubes previously described in literature^17,69^. 20 μL of O/N culture (grown anaerobically in BHI and TY medium) was mixed into tubes containing 0.4 % BHI and TY agar respectively. Strains were grown aerobically at 37°C for 24 h and the growth inhibition was measured by two independent lab members. Strains were assessed for their growth at 1% oxygen in a Don Whitely Scientific MA500 VAIN Workstation, O/N cultures were diluted to an OD_600_ of 0.1 in pre-reduced TY medium and growth was measured every 2 h for 12 h. In parallel, O/N cultures were diluted to an OD_600_ of 1.0, serially diluted, spotted on pre-reduced TY agar and incubated for 24 h.

### Acidic Stress Assay

O/N cultures were diluted to an OD_600_ of 0.1 in pre-reduced BHI-S adjusted to pH 4, 5, 6 and 7. Bacterial growth was measured every 2 h in a spectrophotometer. Viable cells were enumerated onto pre-reduced BHI-S agar plates at 0, 6, 12 and 24 h.

### Oxidative and Nitrosative Stress Detoxification Assay

To assess oxidative stress, O/N cultures were standardised to a McFarland Standard of 1.0 and plated onto pre-reduced BHI agar plates. Sterile 6 mm blank antibiotic disks (Oxoid, UK) were place on the plate and 10 μL of (1, 2, 4 and 9.8 M) H_2_O_2_ and (2, 4, 6 M) methyl viologen (Sigma-Aldrich, UK) was dispensed onto each separate disk. Inhibition of bacterial growth was measured with zone of inhibitions (mm) after 24 h and 48 h incubation.

A spotted dilution assay was conducted in a similar manner to Kint *et al*., 2017. O/N cultures were diluted in pre-reduced sterile PBS and 10 μL were spotted onto pre-reduced TY media supplemented with 200, 500, 1000 and 1500 μM/mL sodium nitroprusside (SNP). After 24 h incubation, bacterial growth was enumerated via CFU/mL and compared to the control agar plates (no SNP)

### Motility Assay

Motility assays were conducted as described in literature^70^. Briefly, 0.3 and % BHI-S agar was used for swimming and swarming assays respectively. 3 μL of O/N culture were spiked into the centre of the agar and incubated unturned at 37 °C. At 24 h and 48 h, the diameter of each sample was measured for motility.

### Sporulation and Germination Assays

Sporulation assays were conducted as described by Edwards and McBride^71^. Spores were also enumerated by phase contrast microscopy on a DM18 (Leica Microsystem, Germany). 2 μL of each sample (controls and post-treatment) were dispensed onto 1% agar pads. 5 randomly selected representation images were captured and the percentage of spores to total cells was calculated.

### Biofilm Formation

The quantification of biofilm biomass by crystal violet (CV) was done in a similar manner to the protocol described in Dapa et al, 2013. O/N cultures were standardised to an OD_600_ of 0.1 and subcultured till OD_600_of 0.5 in BHI-S + 0.1M Glucose (BHI-SG). The cultures were subsequently diluted to an OD_600nm_ of 0.05 and 1 mL was dispensed into pre-reduced 24 well tissue-culture treated plates (Falcon, USA). Plates were incubated for 24 h or 72 h (parafilm was used to secure the lid and plate to avoid excessive evaporation). At each timepoint, each well was gently washed twice with pre-reduced sterile PBS and allowed dry for 30 min, stained with 1 mL 0.2% filter-sterilised (CV) for 30 min. Excess CV was removed, the wells were washed twice with 1 mL PBS and destained with 1 mL of methanol for 30 min. The destained CV was diluted 1:1, 1:10 and 1:100 in methanol and OD_570_ was measured immediately.

For microscopy, biofilms were grown in Nunc™ LabTek^Tm^ II Chamber Slide™ System (Thermofisher) with 1 mL of diluted culture. After 24 h and 72 h, planktonic media was carefully removed and washed twice with 0.1% saponin (w/v%) and incubated with FilmTracer™ LIVE/DEAD™ Biofilm Viability Kit (Invitrogen, USA) following manufacturer’s instructions. The dyes were washed twice with pre-reduced PBS and the biofilm was fixed with 4% paraformaldehyde (w/v) for 15 min. Samples were washed twice with 500 μL sterile water and imaged using with a Perkin Elmer dual-camera spinning disk confocal microscope. Z-stacks of each biofilm was taken at the recommended excitation/emission spectra with increments of μm. 5 representative images were taken and analysed using Fiji (ImageJ).

### Toxin Assay

O/N cultures in BHI-S were subcultured into fresh pre-reduced BHI-S and grown till early stationary (10 h). Each sample was normalised by OD_600_ and the toxin was quantified by separate detection of *C. difficile* toxin A and B ELISA (tgcBIOMICS, Germany) following manufacturer’s instructions.

### RNA Extraction

O/N cultures were mixed with RNA protect (QIAgen, UK) at a ratio of 1:2 and pelleted at 5000xg for 10 min. Samples were resuspended with 600 μL ice-cold LETS buffer (0.1 M LiCl, 0.01 M Na_2_EDTA 0.01M Tris-Cl (pH 7.4) and 0.2% SDS) into Lysing Matrix B tubes (MP Biomedicals, China) and run on a FastPrep-24 5g (MP Biomedical, China) for a total of 6 cycles: 30 seconds at 6.5 m/s and 180 seconds on ice. The tubes were subsequently centrifuged at the highest speed at 4°C for 10 min and the supernatant was transferred to nuclease-free tubes. TRIzol-Chloroform RNA extraction was performed, 1 mL of TRIzol Reagent (ThermoFisher Scientific, UK) was added per 1 ×10^7^ CFU/mL bacteria and incubated at room temperature (RT) for 5 min. 500 μL of chloroform was added, further incubated at RT for 10 min and centrifuged at 16000 xg at 4 °C for 15 min. The supernatant was dispensed into a new nuclease-free tube and 500 μL of isopropanol was added and incubated at RT for 10 min. Samples were centrifuged at 12000 xg at 4 °C for 10 min, the pellet was washed with fresh 70% ethanol and resuspended with nuclease-free water. TURBO DNase (Invitrogen, UK) was used to removed contaminating genomic DNA according to the manufacturer’s protocol. Treated samples were cleaned with 2.5 mM nuclease-free LiCL (Invitrogen, UK) according to manufacturer’s protocol. The purity of each sample was analysed with the RNA Pico 6000 Assay protocol on an Agilent 2000 Bioanalyser, according to manufacturer’s protocol (Agilent, USA).

### RT-qPCR

cDNA was generated from RNA using SuperScript IV Reverse Transcriptase (Invitrogen, USA) following manufacturer’s instructions. Each gene specific primer was designed with Primer-Blast to create a 145-155 bp amplicon with an annealing temperature of approximately 60 °C (Table S2). Samples were amplified with Luna Universal qPCR Master Mix (NEB, USA) according to manufacturer’s instructions in a Mx3005 qPCR system (Agilent, USA). The quantity of cDNA in each reaction was normalised to *gyrA* and relative gene expression was calculated by the 2^-ΔΔCT^ method.

### RNA Sequencing and Analysis

RNA samples were sent to Novogene Ltd for rRNA removal and cDNA sequencing on the Illumina NovaSeq 6000 platform. 150 bp paired end sequences were analysed with FastQC, with approximately 10 million reads per sample. Reads were mapped to *C. difficile* R20291 reference genome (NC_013316.1) with Bowtie2 (version 2.4.5)^72^ and read counts generated using Samtools (1.13) and Bedtools (2.30.0)^73^. DESeq2 was used to calculate differential gene expression using a negative bionomial distribution^74^. Genes were deemed significant when the log_2_ fold-change was ≥ −2 or ≤ 2 and the P adjusted value was ≤ 0.05. Results were visualised with R packages ggplot2 (https://cran.r-project.org/web/packages/ggplot2/index.html) and pheatmaps (https://cran.r-project.org/web/packages/pheatmap/index.html). All sequencing reads were deposited to the European Bioinformatics Institute (Accession number E-MTAB-12114).

### *In vitro* Gut model infection assay

To assess bacterial adhesion to epithelial cells, the experiment was carried as previously described^34^. O/N cultures of bacterial strains grown in BHI-S, standardised to an OD_600_ of 1.0 in DMEM and incubated at 37°C in anaerobic conditions for 1 h. The inoculum was enumerated via CFU/mL. The Snapwell inserts, containing polarised Caco-2, HT29 and CCD 18co myofibroblasts were each sandwiched between two compartments of the VDC (Harvard Apparatus, UK). 3.5 mL of media of pre-warmed DMEM with 10% fetal calf serum (DMEM-10) was added to the basolateral compartment, whilst the apical compartment was infected with the diluted, acclimatised bacterial strains at an MOI of 100:1. Aerobic and anaerobic gas was pumped into each respective compartment at approximately 25 psi, with 15-20 cc/mm per half-cell. The bacterial culture on the apical compartment was washed with PBS and replaced with pre-reduced DMEM-10 at 3 h post-infection. At each desired timepoint, the Snapwells with adhered bacteria were washed twice with pre-reduced 1 mL PBS and lysed with pre-reduced 1 mL sterile water in anaerobic conditions. Adhesion was quantified through serial dilution of Snapwell lysates onto BHI-S agar.

### *Galleria mellonella* infection model for virulence

Galleria mellonella (Livefoods, UK) were stored at 4°C and used within 1 week. Larvae were filtered by skin blemishes, ability to self-upright, weighed (0.25-0.30 g) and swab sterilised with 70% ethanol. O/N cultures were subcultured in pre-reduced BHI-S and allowed to grow till exponential phase, approximately OD_600nm_ of 0.2-0.3. Cultures were centrifuged at 2400 xg at 4°C for 5 min and resuspended in ice-cold BHI-S to limit bacterial growth. 10 μL was administered to each treatment group (of 8 larvae) via oral inoculation between the mandibles using long gel loading tips (Eppendorf, UK). At each desired timepoint, survival rate was numerated through visual observation based on mellonisation and larval movement, with black/brown, immobile insects constituting for death^75^. Larvae were placed on ice (to limit movement) and dissected dorso-ventrally with scissors to obtain gut contents. Extracted guts were placed in 100 μL ice-cold PBS, vortexed at 3000 RPM for 1 min and enumerated on BHI-S agar.

### Immunoblot Assay

*C. difficile* strains were subcultured in BHI-S, TY and TY supplemented with 25 μM/mL SNP for 5 and 10 h, in a similar manner as described previously. 10 mL of each sample was placed on ice and centrifuged at 5000 g for 5 min at 4°C. The supernatants were discarded and the pellets were washed with ice-cold PBS. Bacterial cells were lysed with a freeze-thaw cycle^68^, the pellets were subsequently resuspended in ice-cold PBS with 10 μL/mL Halt™ Protease Inhibitor Cocktail (ThermoFisher, USA) according to manufacturer’s instructions. After incubation at 37°C for 1 h, the pellet was washed and resuspended in sterile water and the protein concentration was quantified by Qubit (ThermoFisher, USA). Samples were normalised to a total concentration of 10 ng, loaded onto a 10% Mini-PROTEAN TGX Stain-Free Precast Gel (BioRad, USA) and transferred onto PVDF using standard methods. The membrane were blocked with 5% bovine serum albumin (Merck Millipore, USA) in TBST (1x Tris-Buffered Saline, 0.1% Tween20 (v/v)), probed with affinity purified anti-σ^B 23^ diluted 1:500 in BSA O/N, followed by anti-rabbit-HRP-linked secondary antibody (Cell Signalling Technology, USA). Blots were stripped using Stripping Buffer (0.2 M glycine, 0.1% SDS (w/v), 1% Tween20 (v/v) and adjusted to pH 2.2), washed thrice, blocked O/N with 5% BSA and probed with anti-R20291 sera.

Crude protein lysates stained with Coomassie Brilliant Blue R-250 (BioRad, USA) were sent for mass spectroscopy at the Warwick Proteomics Research Technology Platform. Gels were digested^76^ and processed with an Orbittrap Fusion with UltiMate 3000 RSLCnano System (Thermo Scientific).

### Statistical Analysis

All data were subjected to the Anderson-Darling and Shapiro-Wilk normality test (dependent on the sample size). To assess significance between two treatment groups, two-tailed Students t-test and Mann-Whitney-Wilcoxon test were used. The significance of percentage survival was examined with the Kaplan-Meier estimator. Significance was denoted by asterisks, with * = *p-value* ≤ 0.05, ** = *p-value* ≤ 0.01, *** = *p-value* ≤ 0.001 and **** = *p-value* ≤ 0.0001. No significance is denoted with the abbreviation *n*.*s*.

## Results

### Deficiency in RsbW and RsbW overexpression do not cause a fitness defect

A defective RsbW protein is thought to negatively impact the overall fitness of the bacterium as previously seen in *B. subtilis*^20,25^, as the presence of unbound σ^B^ could lead to unnecessary transcription and subsequent cellular toxicity. A *rsbW* deletion mutant (Δ*rsbW*) was constructed as described in Methods, such that the start codon of σ^B^ was maintained. To assess overall bacterial fitness, growth of the WT, Δ*rsbW*, and a complemented Δ*rsbW* strain, where the *rsbW* gene was expressed episomally, was monitored in three different growth media: BHI-S medium (Figure S2A), TY medium (Figure S2B) and DMEM (Figure S2C). Over the course of 12 h, no excessive growth defects were observed between the three strains; there were no differences between the mean generation time between strains (Figure S2D). This suggests that the absence of RsbW does not incur a significant fitness defect in *C. difficile*.

Conversely, to assess anhydrotetracycline (aTc)-mediated overexpression of RsbW, each strain was serially diluted onto BHI-S agar supplemented with/without thiamphenicol and aTc(Figure S3A)^23^. Δ*rsbW* appeared to have a slight growth defect (Figure S3B). However, a similar defect was seen in the control strains, indicating that the defect was an effect of aTc rather than overexpression of RsbW. Thus, our data indicate that the absence of or increased expression of RsbW did not impact bacterial fitness *in vitro*.

### RsbW impacts *C. difficile* responses to stress

The role of σ^B^ in *C. difficile* stress response has been characterised^17,18,23,29^, however the contribution of RsbW has not been investigated. We first examined the effects of common environmental stresses that *C. difficile* is likely exposed to in the gut, such as oxygen and pH. When *C. difficile* was grown in soft agar tubes in an aerobic environment as previously described in Kint *et al*, only a small difference between the WT and *rsbW* mutant was observed in the zones of growth (Figure S4A, S4B). However, when the strains were cultured in liquid broth with 1% oxygen, a quicker rate of growth was seen for Δ*rsbW* as compared to the WT and the complemented strain (Figure 1A). When strains were cultured in acidic liquid growth medium at pH 5 (Figure 1B), Δ*rsbW* showed better growth compared the WT and complemented strains, although no differences were observed between strains when cultured in growth media at pH 4, pH 6 and pH 7 (Figure S5A). Enumeration of colony forming units (CFU) from these conditions further confirmed better survival of Δ*rsbW* at pH 5.0 (Figure S5B). A quicker generation time was also indicated for Δ*rsbW* for pH 5, 6 and 7 (Table S3). Reactive oxygen species (ROS) and reactive nitrogen species (RNS) produced by immune cells are key to bacterial killing during infection. During CDI, neutrophils and macrophages are recruited to the site of colonisation releasing ROS and RNS. σ^B^ has been shown to detoxify these, likely through the upregulation of reverse-ruberythrins, NADH-rubredoxin-, NO- and nitro-reductases^17,23,29^. To assess the ability of the Δ*rsbW* to detoxify ROS and RNS, detoxification of H_2_O_2_ and superoxide anion O_2^-^_ (from methyl-viologen) was assessed as described previously^17,18,23^. Both the WT and mutant bacterial lawns showed similar zones of inhibition at 1 M H_2_O_2_ (Figure S6A) and 2 M methyl-viologen (data not shown). However, when exposed to higher concentrations, a clear decrease in the zone of inhibition was visible for the Δ*rsbW* grown in presence of 4 M and 6 M methyl-viologen (Figure 1C). H_2_O_2_ at higher concentrations produced no difference (2 M and 4 M) or a slightly decreased zone of inhibition (9.8M) of Δ*rsbW* (Figure S6A).

**Figure 1.**
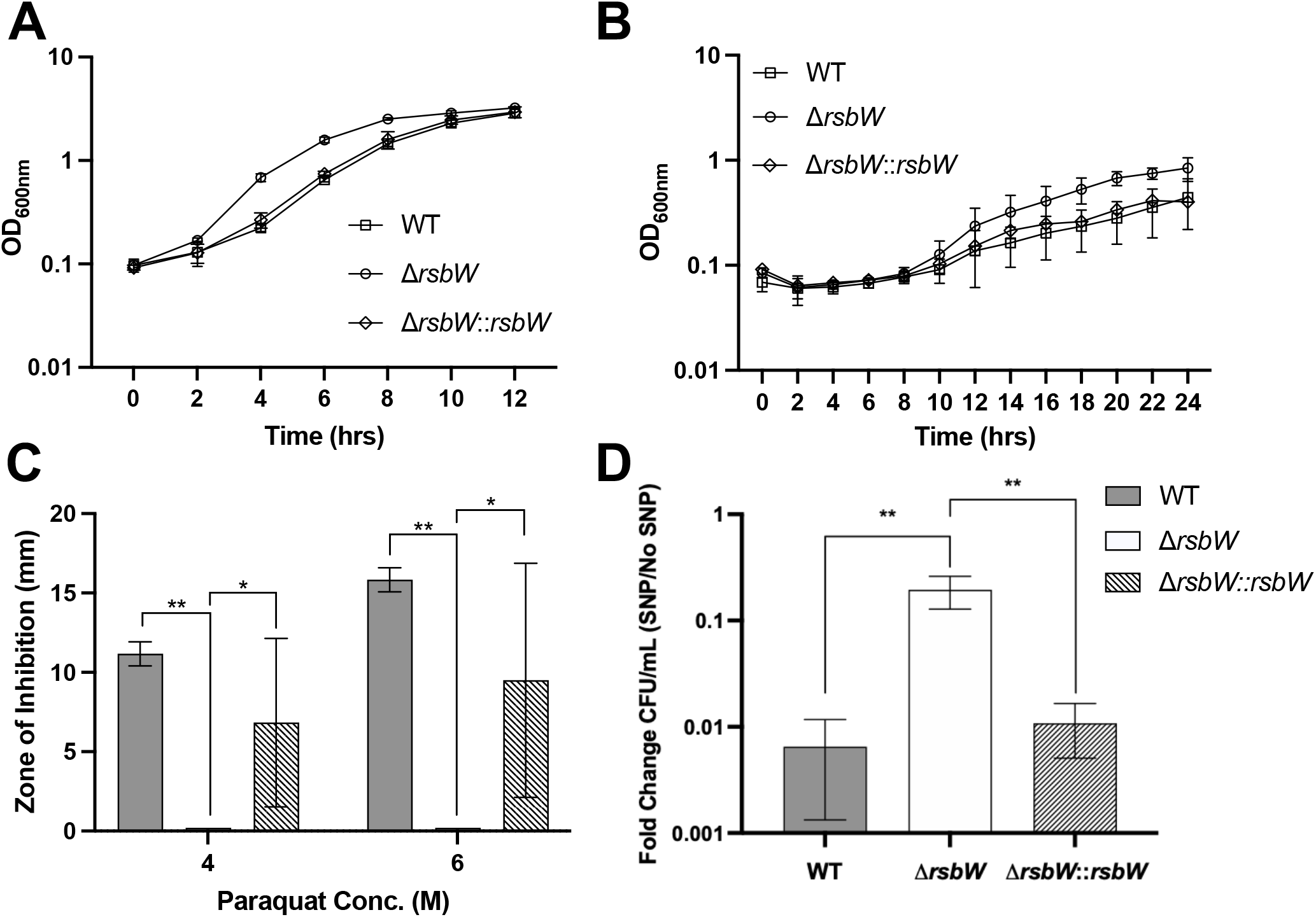
Tolerance of external stresses by Δ*rsbW*. Growth of WT, Δ*rsbW* and complemented strain Δ*rsbW*:*:rsbW* as measured by OD_600_ was compared in A) TY medium in presence of 1% oxygen and B) BHI-S medium adjusted to pH 4. C) The zones of inhibition on BHI plates supplemented with 4 M and 6 M paraquat were compared for the WT, Δ*rsbW* and Δ*rsbW*:*:rsbW*. D) Colony counts for WT, Δ*rsbW*, and Δ*rsbW*:*:rsbW* on BHI-S agar supplemented with and without 1.5 mM sodium nitroprusside. N=3 (with 3 technical replicates/experiment). Error bars indicate standard deviation (SD), significant differences are indicated by * *p-*value < 0.05 and ** *p-*value < 0.01 as determined by Mann-Whitney U test.

As the expression of NO-reductases, nitro-reductases and hydroxylamine reductases are controlled by σ^B^, we next tested sensitivity to RNS^17,18^. Δ*rsbW* growth was not impacted at lower concentrations of 0.2 mM and 0.5 mM (Figure S6B) sodium nitroprusside (SNP), a nitric oxide donor compound, but resulted in a slight difference between the WT (and control strains) and Δ*rsbW* (Figure S6B). However, a significant difference was observed in bacterial survival with SNP at 1.5 mM (Figure 1D), indicating that the Δ*rsbW* is able to reduce nitric oxide (NO) more efficiently than the WT. Thus, these results indicate that the *rsbW* mutant can survive better in conditions of high levels of oxidative and nitrosative stress.

### RsbW controls sporulation

Production of spores is a key mechanism by which *C. difficile* evades unfavourable conditions. Sporulation in *C. difficile* is controlled by many regulators and four sporulation associated σ factors^30,31^. Sporulation is negatively associated with σ^B^; a 10-fold increase in sporulation rate was reported in a σ^B^ mutant while the germination efficiency was unaffected^17^. Our sporulation assays also indicated a similar phenotype, the “always on” σ^B^ in Δ*rsbW* leads to a dramatic decrease in sporulation when examined under phase contrast microscopy (Figure 2A, B). To assess spore viability, the number of spores produced were quantified by measuring CFU in presence of germination agent, sodium taurocholate, after heat or ethanol treatment (Figure 2C). More than a 100-fold difference was observed, with a sporulation frequency of ∼16% for the WT compared to ∼0.05% for Δ*rsbW* in either treatment conditions. The ability to form spores was restored in the *rsbW* complemented strain and a *spo0A* mutant was used as a negative control. Thus, absence of RsbW negatively impacts sporulation.

**Figure 2.**
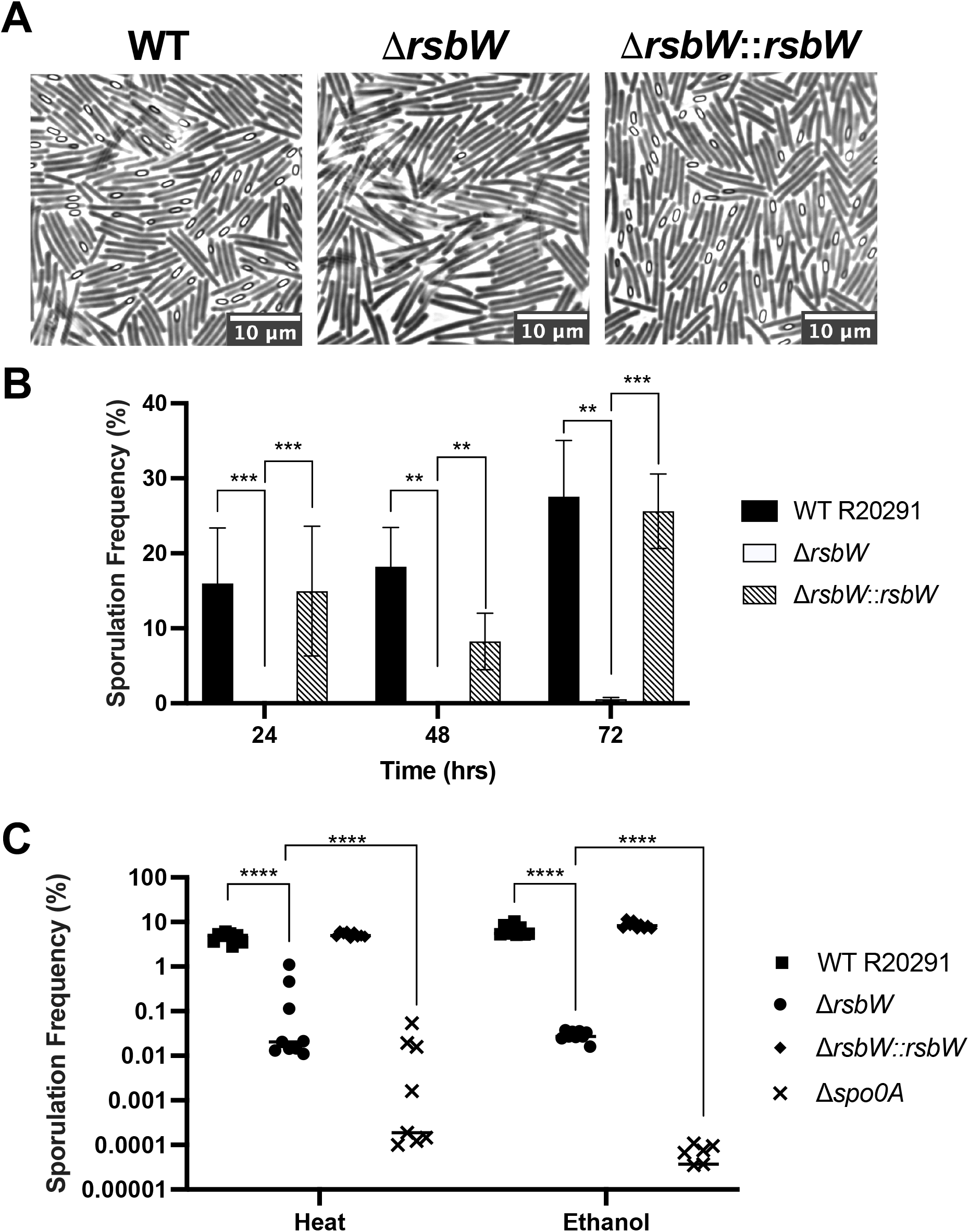
Δ*rsbW* displays severe defects in sporulation. *C. difficile* strains WT, Δ*rsbW* and Δ*rsbW*::*rsbW* were grown on 70:30 sporulation medium and representative phase contrast microscopy images were taken after 24 h (A). Visible spores were enumerated from the microscopy images (N=3, 5 representative fields/ experiment). C) Cultures grown on 70:30 sporulation medium were diluted to OD_600_ 1.0 and subjected to heat and ethanol treatment. Germination frequency was calculated from colony counts of cultures before and after treatment, on BHI-S +/-0.1% taurocholate. N=3 (3 technical replicates/experiment). Error bars indicate SD, significant differences are denoted with ** *p-* value < 0.01, *** *p-*value < 0.001 and **** *p-*value < 0.0001 as determined Student’s t-test or Mann-Whitney U test.

### Biofilm formation is modulated by rsbW

Under unfavourable conditions, *C. difficile* is known to produce a biofilm, which is a dense extracellular matrix that forms a physical barrier to dampen environmental stresses^32,33^. Biofilms formed by WT and Δ*rsbW* were quantified using crystal violet staining at 24 h and 72 h. At 24 h (Figure 3A), no significant difference was observed between the strains, although at 72 h, Δ*rsbW* appeared to produce more biomass overall compared to the WT (*p*-value = 0.008).

**Figure 3.**
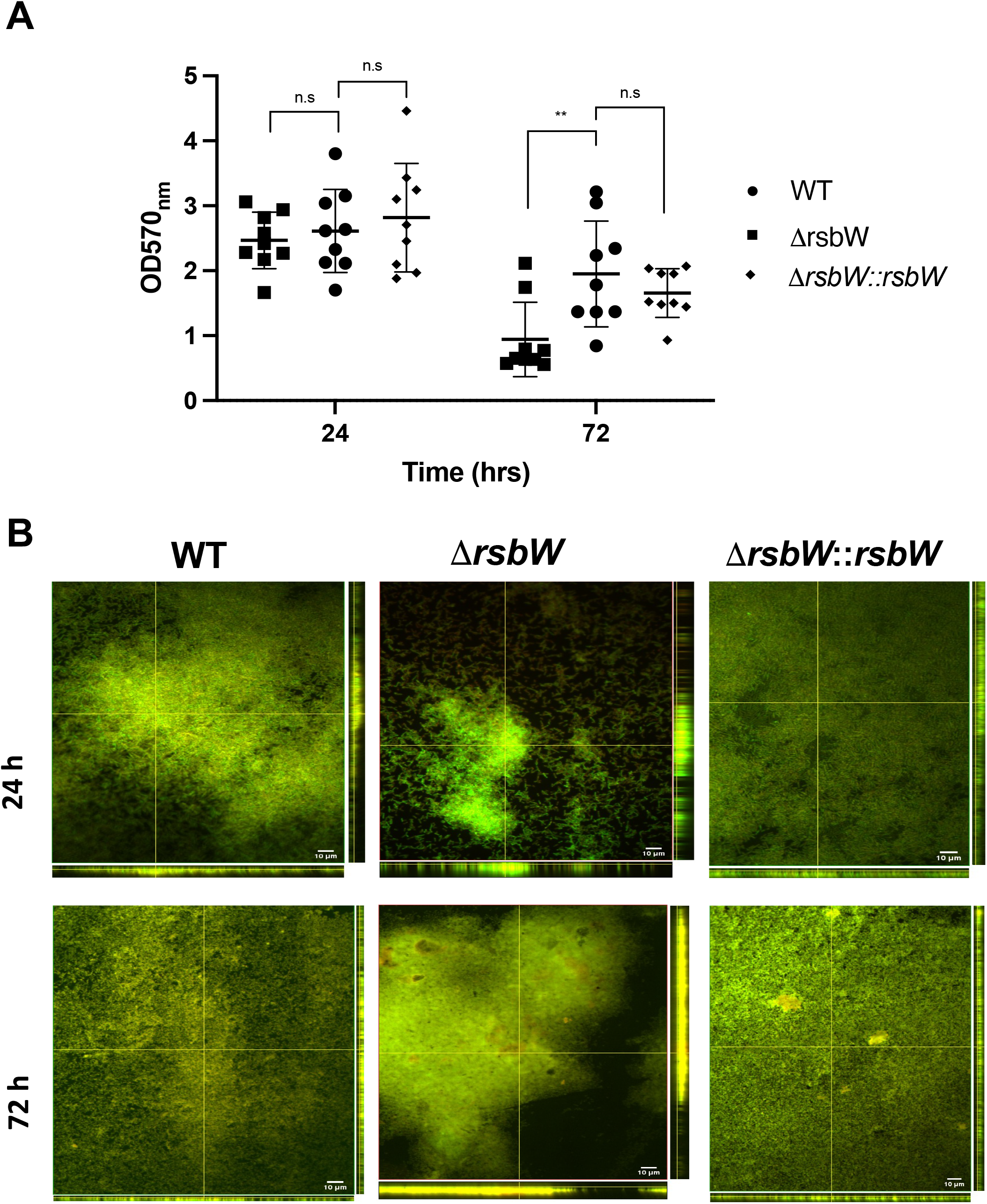
Biofilm formation is modulated by RsbW. *C. difficile* strains were grown in BHI-S +0.1 M glucose during early exponential phase for 24 h and 72 h. A) Biofilm biomass was quantified by crystal violet at OD_600nm_ in 24-well tissue-culture treated polystyrene plates N=3 (3 technical replicates/experiment). B) 24 h and 72 h biofilms were stained with FilmTracer™ LIVE/DEAD®. Orthogonal views of the biofilm z-stack depict biofilm thickness. N=3, 5 representative images/experiment. Error bars indicate SD, no significance was denoted with n.s and significance with ** *p-*value < 0.01 as determined Student’s t-test or Mann-Whitney U test.

To probe these differences further, biofilms were stained with a LIVE/DEAD and examined by confocal microscopy. Biofilms formed by the WT and Δ*rsbW* strain exhibited different characteristics (Figure 3B) with the WT forming consistent, thin biofilms at both 24 h and 72 h and Δ*rsbW* forming a more varied, thicker, but sparser biofilms. The complemented strain behaved in a similar manner to the WT strain, however some thick biofilm structures were intermittently spotted (Figure S7) and showed similar distribution to Figure 3A. Thus, RsbW may play a role in modulating *C. difficile* biofilm formation. The increased differences seen at 72 h could suggest the role of σ^B^ in a nutritional stress response due to a gradual depletion of nutrients in the growth medium.

### Δ*rsbW* adheres to epithelial cells better in a human *in vitro* gut model

The gut mucosa provides invading bacteria with a range of environmental insults. σ^B^ has been associated with increased colonisation in a dixenic mouse infection model^17^, however its interactions with human epithelial cells have not been studied. Using an *in vitro* gut model, previously used to probe for *C. difficile* bacterial adhesion^34^, we compared the attachment of WT and Δ*rsbW* to a multi cellular-layer of Caco-2, HT-29 MTX E12 and CCD-18co cells in a vertical diffusion chamber (VDC). This dual compartment system allows bacterial growth in anaerobic conditions and keeps the epithelial cells in an oxygenated environment. VDCs were infected with WT or Δ*rsbW* at an MOI of 100:1 for 3 h, 6 h and 24 h. Non-associated bacteria were washed off and the adherent bacteria were enumerated via CFU. Across each timepoint, more Δ*rsbW* were recovered from the monolayer compared to the WT (Figure 4A). At 3 h, Δ*rsbW* was able to adhere ∼2-fold more than the WT, increasing to an average 4-fold difference at 6 h. This increased difference at 6 h may suggest that Δ*rsbW* can tolerate host-mediated stress better to outgrow the parental strain. Although more variation was observed at 24 h, a similar average difference of 3.5-fold was observed between the WT and mutant strain.

**Figure 4.**
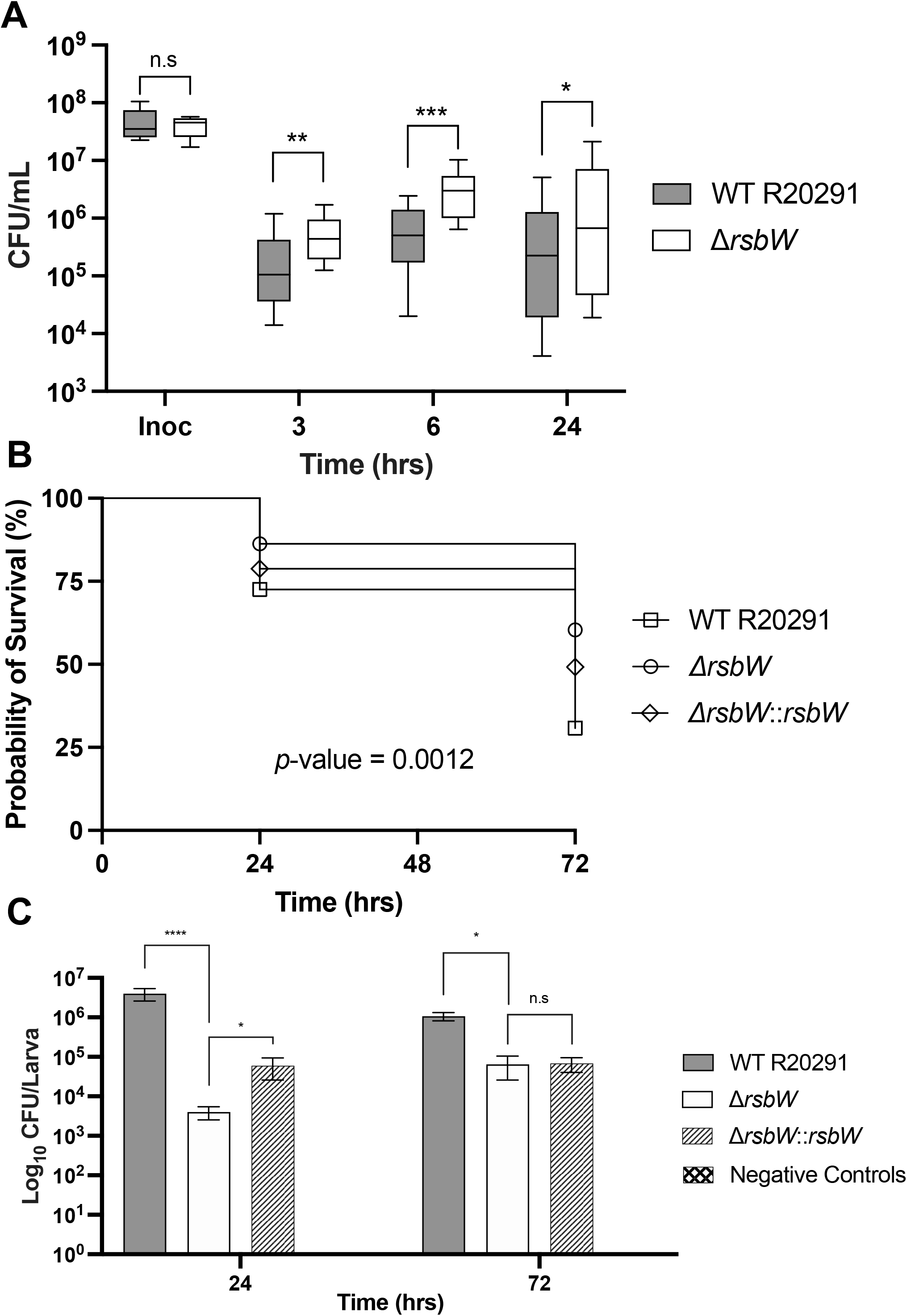
Δ*rsbW* demonstrates increased cell adhesion but decreased virulence. A) WT *C. difficile* and Δ*rsbW* adherence to epithelial cells across 3 h, 6 h, 12 h and 24 h after infection in an *in vitro* VDC infection model N=3 (3 technical replicates/experiment). B) Survival curve of *Galleria mellonella* infected with *C. difficile* WT, Δ*rsbW* and Δ*rsbW*::*rsbW* strains across 24 h and 72 h (N=5, 8 larvae/strain/timepoint) C) Adherent bacterial population recovered from infected *G. mellonella* gut enumerated with CFU/mL on BHI-S agar. Error bars indicate SD, no significance was denoted with n.s and significance with * *p-*value < 0.05, \*\**p-*value < 0.01, *** *p-*value < 0.001, **** *p-*value < 0.0001 as determined by Mann-Whitney U test and Log Rank Mantel Cox test for the survival curve.

### RsbW impacts colonisation in a Galleria infection model

An insect infection model was used to study the role of RsbW in colonisation. *G. mellonella*, previously used to assess the efficacy of phage therapy in treating *C. difficile* infection^35,36^, is noted for ease of use and cost-efficiency. We infected ethanol-sterilised larvae with approximately 1 ×10^5^ CFU of WT, Δ*rsbW* or *rsbW* complemented strain through oral gavage and the infection outcome was monitored over 72 h. Any larvae that displayed immobility, melanization (brown and hardened blemishes) or had turned black were deemed deceased^36^. Figure 4B, shows *G. mellonella* infected with Δ*rsbW* displayed a higher survival rate in both 24 h and 72 h, compared to the WT strain. At 24 h, larvae infected with WT R20291 had a survival rate of 73%, while Δ*rsbW* had 86%, but at 72h, 70% of the larvae had succumbed to infection with the WT strain, as opposed to 40% when infected with Δ*rsbW*. The difference between the two strains was significant with a *p*-value of 0.001 using the log rank test. The complemented strain showed a survival rate more akin to the WT strain. At each time point, the gut from each larva was extracted and CFU enumerated (Figure 4C). The bacterial numbers show a similar trend to the survival assay (Figure 4B): at 24 h a 3-log fold difference was observed between both strains. At 72 h, the difference between recovered bacterial loads decreased by 1-log fold. While we report a regulatory role for RsbW in adherence, we see a decreased colonisation of insect guts, indicating that while bacteria that lack RsbW could attach to the gut surface, perhaps they are not able to survive the complex stresses present in this environment.

### Expression of σ^B^ associated genes in Δ*rsbW*

As some of the phenotypes observed for the *rsbW* mutant were distinct to those reported for the σ^B^ mutant we wanted to understand the effect of *rsbW* deletion on the σ^B^-modulated genes. The transcriptomic profiles of WT R20291 and Δ*rsbW* strains in early stationary phase (10h) were compared by RNA sequencing (RNASeq). Biological replicates of WT andΔ*rsbW* were each seen to cluster together in PCA plots (Fig S8A). Correlation analysis also showed low variation between samples (Fig S8B). In total, 234 and 483 genes were up- and down-regulated respectively (Figure 5A) in the Δ*rsbW* compared to the WT strain. The complete list of differentially regulated genes is included in Table S4. Surprisingly, majority of σ^B^-controlled genes were unchanged or down-regulated as opposed to what was previous reported in the σ^B^ mutants^17,23^. This discrepancy was also observed with genes responsible for oxygen tolerance, oxidative and nitrosative stress responses and acid tolerance. Among the 200 genes thought be controlled by four sigma factors^30^, 71 genes have been associated directly with sporulation. Only 17 sporulation genes were significantly downregulated (Figure 5C), with an emphasis towards spore coat proteins (*cotE, BclA2, BclA3*, CDR20291_RS03125 and CDR20291_RS01475)^37,38^ and mid-to late-stage sporulation genes (*spoIVB, spoVAE, spoIIIAA, spoVAD, spoVAC*). This transcriptional change to prevent spore maturation and release can be reflected in the sporulation frequency observed in Figure 4A. While the global regulator Spo0A associated with spore formation ^32^ was unchanged, interestingly both genes of the SinRR’ locus were dramatically upregulated by ∼5-fold each. The SinRR’ locus encodes a pleiotropic regulator, which is also responsible for sporulation, motility and toxin expression^39,40^.

**Figure 5.**
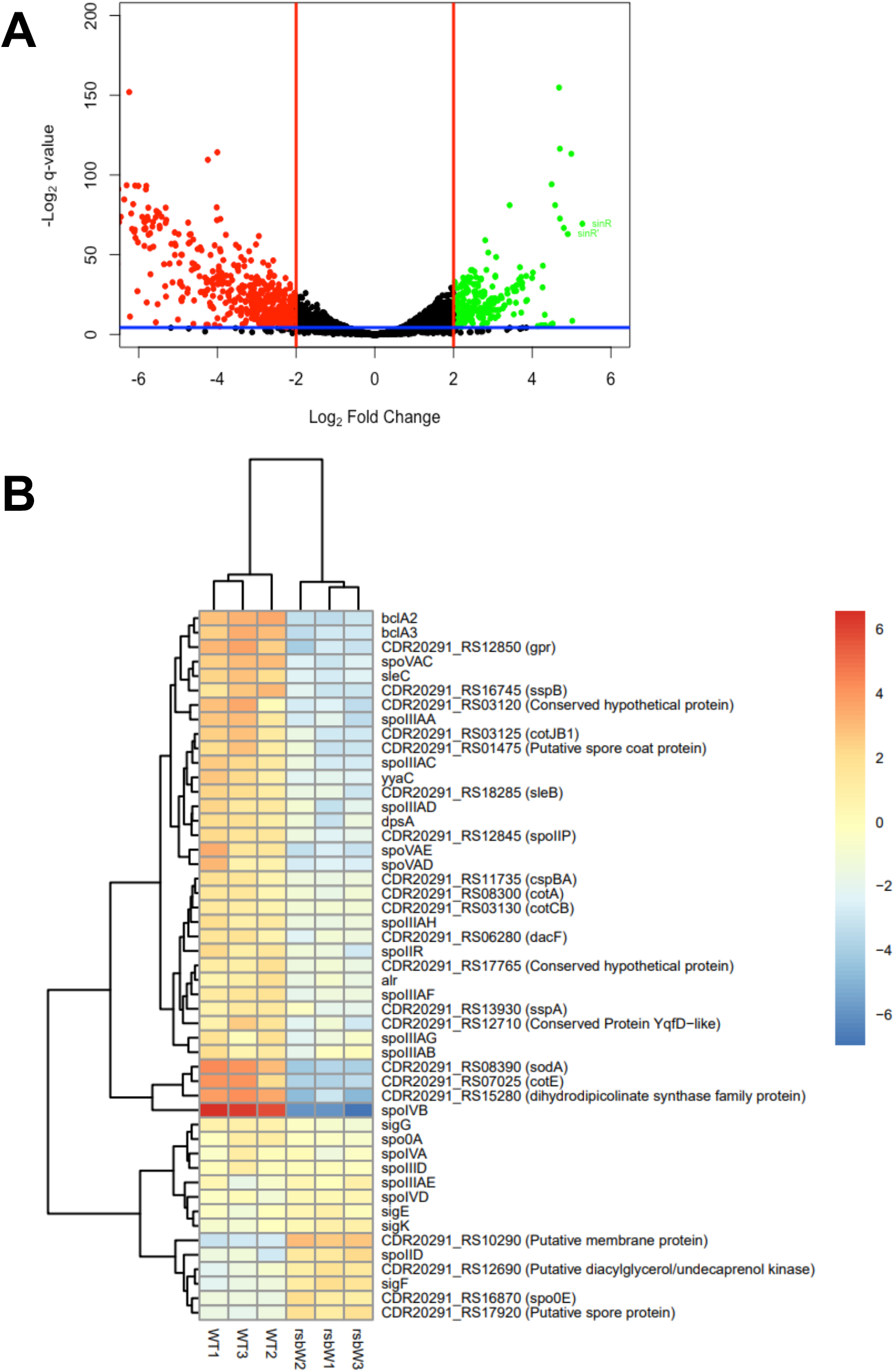
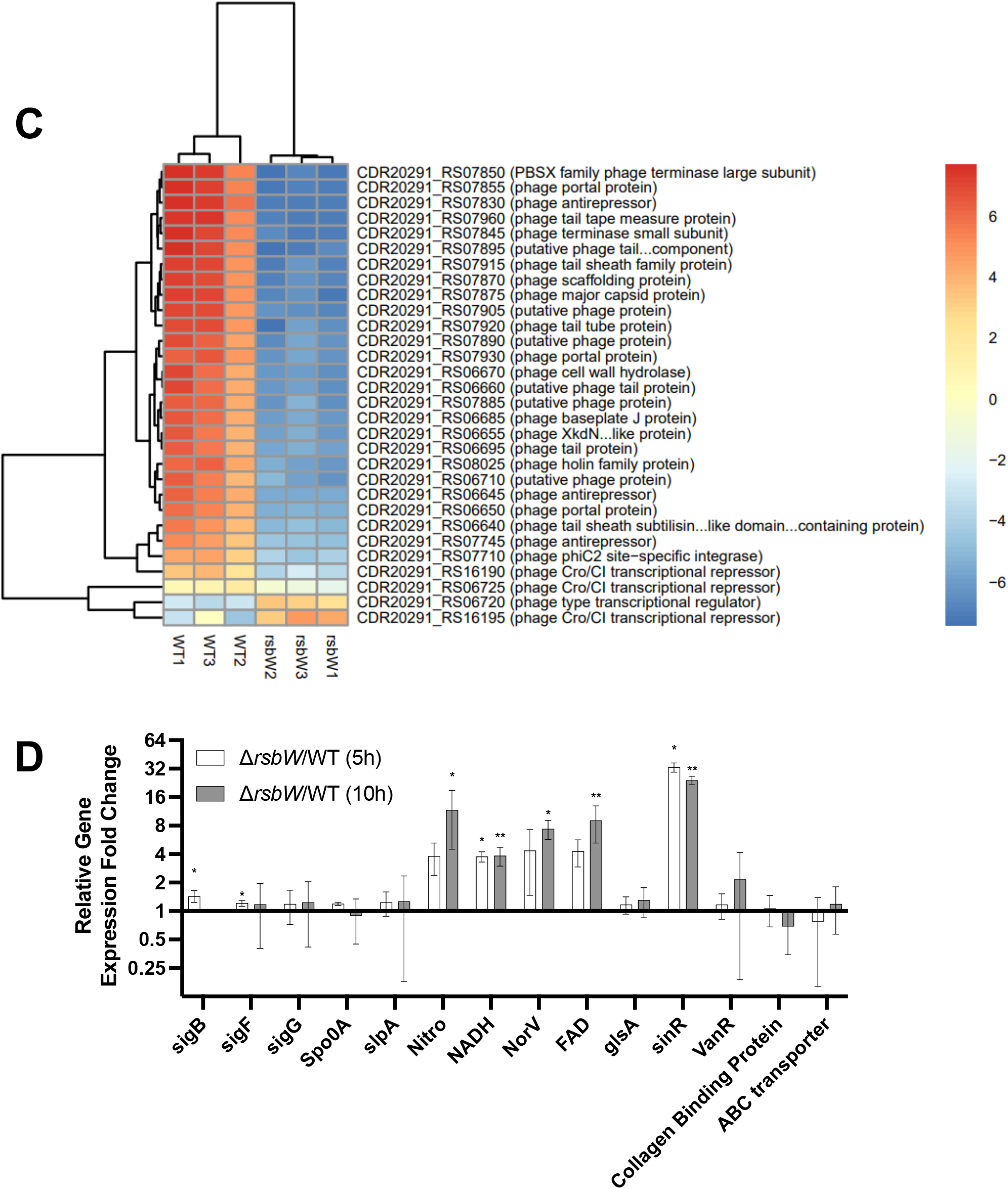
Bacterial transcriptomics of Δ*rsbW* reveals upregulation of the SinRR’ locus. A) Volcano plot reveals differential gene expression when early stationary phase (10 h) Δ*rsbW* is compared to the WT. N=3, *p-*value < 0.05 Heatmap representation of B) sporulation genes and C) phage-associated genes that were differentially expressed in Δ*rsbW*, blue and red gradients indicate down- and up-regulation respectively when compared to WT. D) RT-qPCR confirms the differential expression of selected genes at 5 h and 10 h. Error bars indicate SD, significance was denoted with * *p-*value < 0.05 and \*\**p-*value < 0.01 using a one sample t-test.

Flagella and pilin genes were also downregulated, however, no difference was seen between strains in motility assays of 0.3% and 0.4% agar (Figure S9A and S9B respectively). Interestingly, *tcdA* and *tcdB* were downregulated by 2-fold and 0.5-fold respectively, the expression of toxins through σ^B^ has not been described and could suggest the involvement of another regulator. Toxins A and B measurements from the WT and Δ*rsbW* planktonic cultures indicated a decrease in toxin production, although the differences were not statistically significant (Figure S9C).

In terms of genes associated with biofilm formation, no significant genes were up- or downregulated, with the exception of c-di-GMP-independent VaFE repeat-containing surface-anchored protein (CDR20291_RS18550) which was downregulated 2.6 fold^41^. Other surface proteins CDR20291_RS14700 and *PilA1* was downregulated ∼1.5 fold whilst C40 peptidase surface protein was upregulated approximately ∼1 fold^41,42^. CDR20291_RS14700 encodes for a c-di-GMP controlled surface protein, likely to be controlled by diguanylate cyclase (*DccA*) and phosphodiesterase (*PdcA*) were upregulated by ∼1-fold^42,43^. No distinctive changes were observed in gene expression for cell wall proteins or adhesins, except for Cwp27 which has yet to be characterised^44^. There was no significant change in the expression of genes encoding RsbW and σ^B^. Due to the overlapping stop-start codon, small transcripts of *rsbW* were counted, however manual inspection of reads in Artemis confirmed a *rsbW* deletion in the mutant.

Finally, it was interesting to see differential expression of many phage-associated genes (*p*adj <0.05) in Δ*rsbW*. The majority of phage-associated genes were from incomplete and intact phages ΦMMP04 and ΦC2, most of which were down-regulated. The genes belonging to ΦMMP04 (Figure 5C) are either unchanged or severely downregulated, possibly controlled by CDR20291_RS06720, a CI repressor. A similar transcriptomic profile can be observed in ΦC2 (data not shown); however, the role of the Cro-CI bistable switch is less clear at this instance. Expression of selected differentially expressed genes were further confirmed by qRT-PCR. Most genes showed similar changes in expression, (Figure 5D), aside from genes thought to be controlled by σ^B^ (Nitroreductase, NADH peroxidase, NorV and FAD-dependent oxidoreductase).

Thus, the RNAseq studies indicate that the absence of rsbW is associated with transcriptional profiles that are distinct to the σ^B^ regulon, which may in part be attributed to the induction of other transcriptional regulators like SinRR’.

### Intracellular concentrations of σ^B^ are lower in unstressed Δ*rsbW*

To understand phenotypes displayed by Δ*rsbW* and its distinct transcriptomic profile, the intracellular concentrations of σ^B^ were compared between WT and Δ*rsbW* (anti-σ^B^ antibody kindly provided by Dr W. K. Smits, Leiden University Medical Centre). The relative quantities of σ^B^ were measured in planktonic cultures during exponential and early stationary phase (5 h and 10 h) and in non-stressed and stressed conditions (induced by NO). Surprisingly, during non-stressed growth conditions, a lower concentration of σ^B^ was detected in Δ*rsbW* compared to the WT for exponentially growing bacteria (Figure 6A). Conversely, when the bacteria are exposed to nitrosative stress, similar levels of σ^B^ was observed in both strains (Figure 6A and B). This difference was not observed in the same cultures at early stationary phase (10 h) as intracellular concentrations of σ^B^ remained consistently lower in the mutant strain (Figure 6C). Complete phenotypic complementation was difficult to achieve, as artificial overexpression of σ^B^ was possibly counteracted by this undefined degradation mechanism. Furthermore, the SDS-PAGE protein profiles of total bacteria lysates revealed an altered abundance of proteins for Δ*rsbW* compared to the WT (Figure S10A) in both 5 h and 10 h. Mass spectrometry of the bands that were prominently seen in the *rsbW* mutant showed a higher relative amount of two reverse rubrerythrins (CDR20291_RS07230 and CDR20291_RS07495) in Δ*rsbW* (Figure S10B and S10C), which are known to be controlled by a σ^B^ promoter^29^.

**Figure 6.**
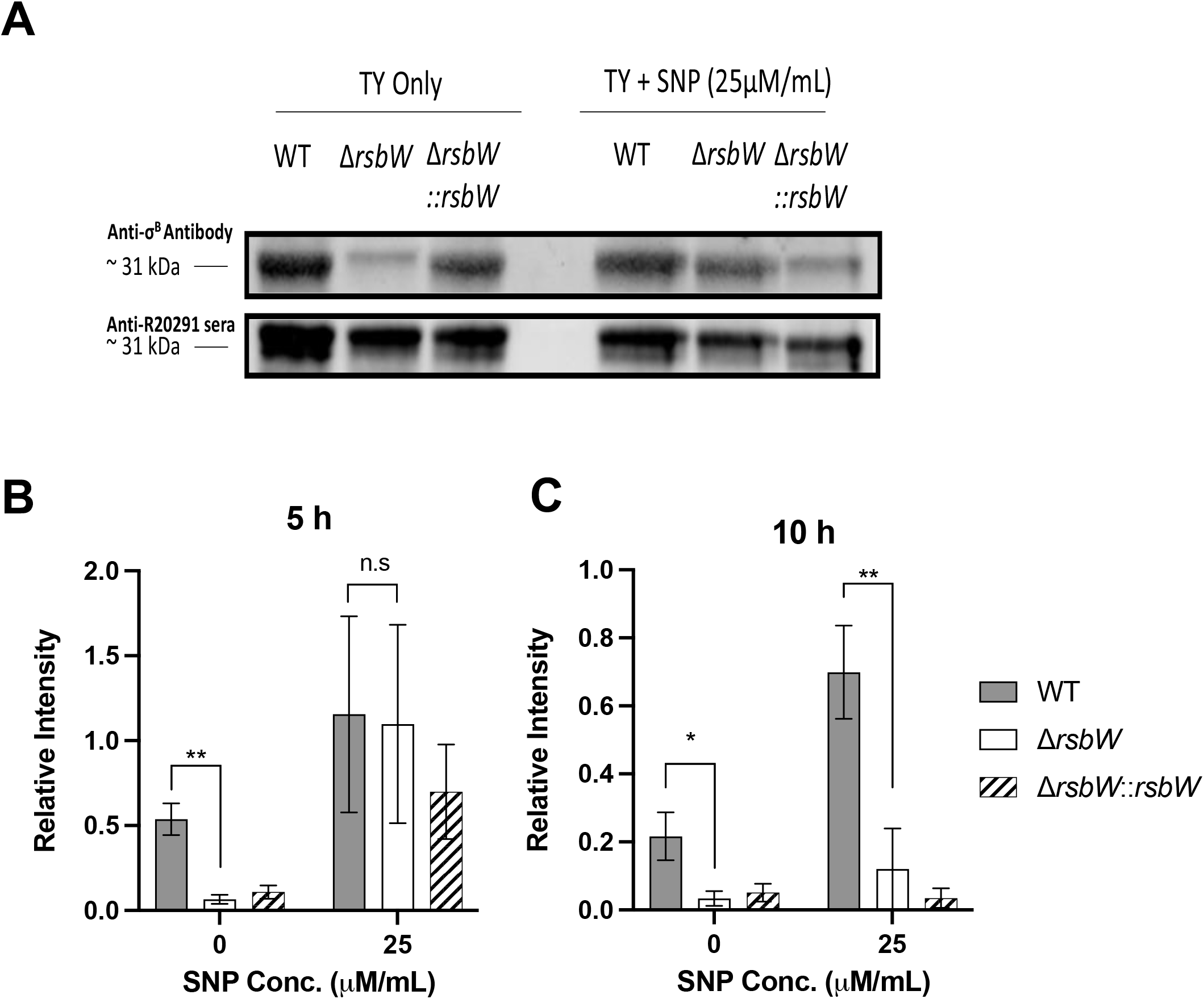
Immunoblotting shows altered relative concentrations of intracellular σ^B^ in Δ*rsbW*. Bacterial whole cell lysates from *C. difficile* strains *C. difficile* WT, Δ*rsbW* and Δ*rsbW*::*rsbW* grown in TY broth +/-SNP to early exponential phase (5 h) or early stationary phase (10 h) were analysed by immunoblotting with anti σ^B^ or anti-R20291 sera (for normalisation) Image representative of 3 independent experiments **(A)**. Band intensities were calculated using Fiji (ImageJ) to determine relative intracellular σ^B^ concentrations at 5 h **(B)** and 10 h **(C)** +/-25 µM SNP. Error bars indicate SD, no significance was denoted with n.s and significance with \**p-* value < 0.05 and \*\**p-*value < 0.01 as determined by Student’s t-test.

Thus, our data indicates that in absence of stress, σ^B^ is likely removed or degraded in the absence of RsbW by an unknown σ^B^ degradation mechanism, which may explain the lack of growth aberration in Δ*rsbW*. Under conditions of nitrosative stress, σ^B^ is not removed or degraded, indicating the necessity of σ^B^ in RNS detoxification.

## Discussion

This study presents a unique insight into σ^B^ activation and control through the anti-sigma factor RsbW in *C. difficile*, based on findings from deletion mutant of *rsbW*. We report here that RsbW modulates many σ^B^-mediated stress responses, as evident from many stress-associated and virulence phenotypes. However, several genes which have not been typically associated with σ^B^, including the SinRR’ pleiotropic regulator are also controlled directly or indirectly by RsbW, indicating additional layers of regulatory control. Furthermore, our data also suggest stress-dependent post-translational control of σ^B^ levels in *C. difficile*.

To our knowledge this is the first description of a targeted *rsbW* bacterial mutant. Previous studies have attempted to create isogenic *rsbW* mutants in other bacteria: in depth studies in Bacillus have not been successful due to a toxic phenotype^20,25^. In *Staphylococcus aureus*, where RsbW and σ^B^ are expressed through coupled translation, the *rsbW* mutant showed a σ^B^ mutant phenotype^45^. Unlike other Firmicutes, the σ^B^ operon in *C. difficile* is under basic control by the housekeeping σ^A^, without a σ^B^ promoter and hence devoid of a positive feedback loop. Thus, we would expect the absence of RsbW in *C. difficile* would lead to unbound σB, in an ‘always on’ manner, without a fitness defect.

While as expected the Δ*rsbW* did not have a fitness defect in normal growth conditions, it was able to respond quicker to low oxygen concentrations, detoxify ROS/RNS stress and acidic environments and thus survive better in these conditions. This can be observed in growth curves in 1% oxygen, 25 µM/mL SNP and pH 5. However, more extreme conditions were required to elicit a difference, as the WT strain was able to tolerate higher concentrations than described in literature^17^. Therefore, semi-lethal stress conditions can crudely distinguish the presence/absence of the partner-switching mechanism. The differences seen between ROS inducers H_2_O_2_ and methyl-viologen may be due to the requirement of an additional enzyme to neutralise by the latter. In the detoxification of methyl-viologen, *B. subtilis* employs superoxide dismutase (*sodA*), which catalyzes the disproportionation of O_2^-^_ into O_2_ and H_2_O_2 ^46^_. The WT and complement strains likely succumb to the effects of superoxide before the stress response genes are activated. However, *sodA* in *C. difficile* has only been associated with spore coat formation or maturation^37^ and not in aerobic stress^47,48^. Instead σ^B^-associated desulfoferrodoxin, reverse rubrerythrins and NADH-rubredoxin reductases have been described as the remedial agent^17^. In line with this, some reverse rubrerythrins were produced at high levels in Δ*rsbW*, as shown by mass spectrometry. The availability of constitutively produced reverse rubrerythrins allows the mutant to tolerate low oxygen and oxidative stress better and promotes enables quicker growth. Although transcriptomic analysis indicates a mild downregulation or no significant difference in the expression of the two reverse rubrerythins, an upregulation of one of these (NADH peroxidase) was seen in the RT-qPCR assay.

Sporulation and biofilm formation, both phenomena that help the bacterium persist under conditions of stress, were impacted in the absence of RsbW. Sporulation was clearly decreased in the absence of RsbW, and this was supported by the transcriptional analyses which showed downregulation of late-stage sporulation genes which are involved in the maturation and release of spores. This aligns with the increased sporulation rate reported for *C. difficile* σ^B^ mutants^17,49^. An increased biofilm formation was observed in the absence of RsbW and the differences in biofilm formation were maximal at later stages of biofilm growth, indicating a role for nutritional stress in this phenotype. A defect in the ability to form biofilms was not seen with *C. difficile* σ^B^ mutants^17^, although σ^B^ has been associated with biofilm formation in *Staphylococcus aureus*, which controls fibronectin-binding protein A expression^50,51^. *C. difficile* encodes orthologues of fibronectin-binding proteins, with FbpA demonstrated to have a role in colonization^52^. FbpA has also been associated with early biofilm formation in the presence of elevated c-di-GMP^41^. Another orthologue, SrtB-anchored collagen-binding adhesin, is also present in *C. difficile* and has a predicted upstream σ^B^ promoter. Although no significant transcriptional changes were observed with any surface proteins associated with *C. difficile* colonization, it is likely that these genes would be expressed only under the relevant conditions of stress, and not during planktonic growth.

Bacteria are exposed different types of stresses, including soluble mediators and antimicrobial peptides, when they are in contact of mammalian cells. We see an increased adhesion with Δ*rsbW* in an *in vitro* gut model used in this study where in addition to cell associated factors, bacteria are likely also exposed to low levels of oxygen and metabolic stress. A 10-fold decrease in colonization was previously reported when σ^B^ was inactivated in 630Δerm, hence we would expect that the Δ*rsbW* under stress is able to associate quicker with the epithelium and multiply earlier. Although it displayed increased adhesive ability *in vitro*, interestingly Δ*rsbW* did not infect as well as the WT in a *G. mellonella* infection model across 24 h and 72 h. The insect gut could be a more complex environment, needing a better regulation of stress responses, which was affected in the absence of RsbW. On the other hand, as noted previously, a difference between the WT and Δ*rsbW* was observed only in extreme stress conditions, which the insect infection model might not be able to provide. Therefore, the lower recovered bacterial numbers could be the result of the partially stressed bacterium possessing a low intracellular concentration of σB.

The transcriptomic profile of Δ*rsbW* unexpectedly revealed the lack of differential expression of several genes in the σ^B^ regulon as previously described^17,23^. Although as expected the transcription σ^B^ gene per se did not change, the lower intracellular concentration of σ^B^ in the Δ*rsbW* in a non-stressed state suggests that the RNAseq profiles may mimic a σ^B^ deletion mutant. Thus, these data may further indicate that exposure to stress is necessary to understand genes truly associated with σ^B^. Indeed, we cannot rule out that RsbW may interact with sigma factors other than σ^B^ and mediate control of additional sets of genes. One of the interesting gene loci that showed a large upregulation in the Δ*rsbW* is the SinRR’ locus; it was upregulated in both 5 h and 10 h planktonic cultures. SinR and SinR’ are transcribed as a single transcript and has not been associated with the control σ^B^ expression, nor has a σ^B^ promoter been identified upstream of the locus. The *sinRR’* expression can be supressed by Spo0A, as a phosphorylated Spo0A (Spo0A∼P) can bind to the only promoter sequence upstream of this locus^53^. While this locus and σ^B^ may contribute to Δ*rsbW* phenotypic responses, SinR regulates the transcription of other pleiotropic regulators such as σ^D^, Spo0A and CodY, which can regulate many other genes in *C. difficile* ^39,54–56^. A direct link between σ^B^ and sinRR’ locus has yet to be described in *C. difficile*. In *B. subtilis*, Spo0A is negatively controlled by Spo0E through dephosphorylation, furthermore Spo0E is positively controlled by a σ^B^ promoter^49^. Additionally, it has been suggested that SinR is able to regulate *spo0A* expression through an unknown mechanism ^53^. The SinR homolog in *B. subtilis* functions as a repressor for the spo0A promoter ^57^. Our transcriptomic data indicates no changes in *spo0A* expression but *spo0E* is but regulated by ∼1.5 fold. Hence, we propose that σ^B^ regulates the *sinRR’* locus (Figure 7) through Spo0E, through dephosphorylation of Spo0A∼P to Spo0A, and subsequent derepression of the sinRR’ locus.

**Figure 7.**
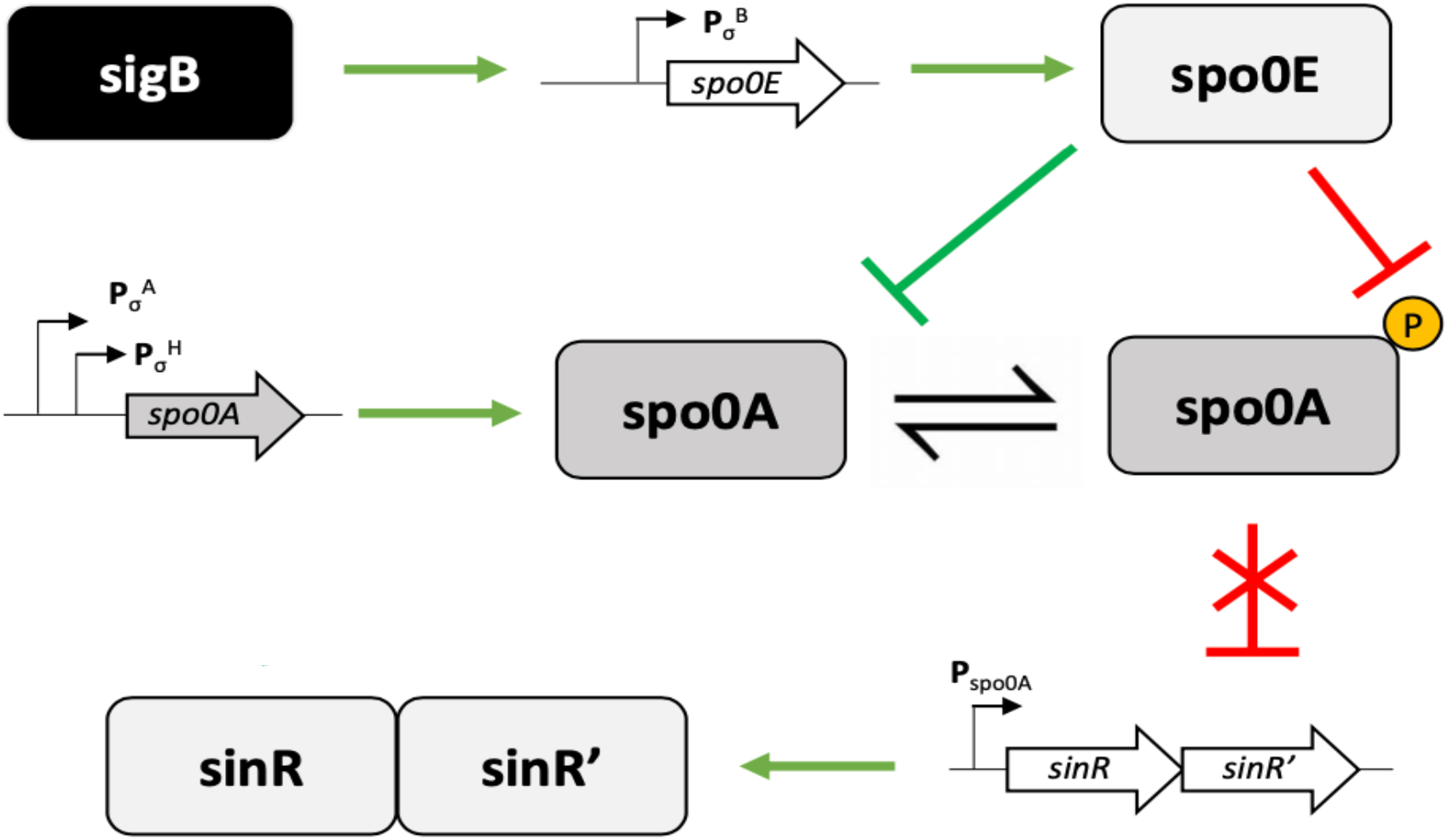
A model for the activation of the *sinRR’* locus through σ^B^. Our proposed model is that σ^B^ activates expression of *spo0E* though a predicted σ^B^ promoter found upstream of *spo0E*. Spo0E dephosphorylates Spo0A∼P to Spo0A, which derepresses the SinRR’ locus. In the absence of RsbW, we have low but constitutive σ^B^ expression which would ultimately leading to an increased sinRR’ expression.

Another intriguing finding was the downregulation of several phage structural and functional genes in the mutant. Previous work from our group has shown that the phage lysis of WT R20291 may contribute to biofilm formation through eDNA accumulation^58^. Yet, the Δ*rsbW* formed better biofilms than the WT R20291 strain. This may suggest that the induced SinRR’ system may play more prominent role the biofilm formation in *C. difficile*. Notably, a downregulation of predicted ‘lytic’ genes was evident. Phaster annotation of R20291 genome describe two incomplete and one complete prophage regions, including ΦMMP04 and ΦC2, which are previously described *Myoviridae* phages^59,60^. However, their association with σ^B^ has not been reported. Lytic genes are under the control of three genes; *int, cro* and *ci*, which control the lytic and lysogenic lifecycle; integrases and Cro control the lytic lifestyle of the prophage, whilst CI (represses *int)* maintains lysogenic stability^61,62^. Cro/CI form a double negative feedback loop. The effect of σ^B^ on both proteins is the clearest in ΦMMP04, where genes encoding CI (CDR20291_RS06720) is upregulated and Cro (CDR20291_RS06725) is downregulated. Although ΦMMP04 does not have an annotated integrase in the prophage region^60^, the majority of its predicted lytic genes are downregulated, presumably as a consequence. In ΦC2, the Cro/CI repressors are not annotated, however a number of helixturn-helix transcriptional regulators, bearing homology to Cro/CI, are differentially expressed. Consequently, *int* (CDR20291_RS07710) and approximately 50 other genes in the prophage region are downregulated. Therefore, σ^B^ could potentially exert control for lysogenic stability, previously associated between σ^H^ and *S. aureus*^63^.

It was surprising that there was a significantly lower concentration of intracellular σ^B^ in Δ*rsbW* in a non-stressed condition, as one would expect that σ^B^ accumulates in the absence of RsbW. However, similar concentrations as the WT were observed when exposed to stress conditions (induced by SNP). A previous study in *B. subtilis*, reported a 10% decrease under non-stressed condition, however the relative concentration of σ^B^ was not measured in a stressed state^19^. A lower concentration of unbound σ^B^ would result in lower or negligible expression of unnecessary genes, and thus not cause any impact on bacterial fitness. This may suggest an unknown proteolytic/ σ^B^ degradation system which is induced during non-stressed states. Caseinolytic proteases (Clp) are one of the main systems employed by bacteria for regulatory protein homeostasis^64^, Clp homologues in Firmicutes and *C. difficile* have been associated with growth and virulence ^65–67^. While it is possible that Clp could control σ^B^, we cannot exclude other posttranscriptional control mechanisms that control translation of σ^B^.

To conclude, we report the unique phenotype of a *C. difficile* strain that constitutively expresses σ^B^ in the absence of the anti-sigma factor RsbW, in stress response, persistence and infection. This study provides new insight into the σ^B^ controlled regulatory circuits in *C. difficile*, including a potential role for σ^B^ in the control of the sinRR’.

## Acknowledgements

J.K.J.C. was funded by a PhD studentship award by the BBSRC Midlands Integrative Biosciences Training Partnership (Grant number 1897785) and a grant from the Warwick Quantitative Biomedicine Programme, We acknowledge the Warwick Computing and Advanced Microscopy Unit for their support & assistance.

J.K.J.C, T.D and R.S made all strains in the study. J.K.J.C, I.Y.L.C and T.M performed the experiments for this study. J.K.J.C, I.Y.L.C and MU were involved in designing experiments in the study. J.K.J.C, I.Y.L.C and MU wrote and reviewed the manuscript. We declare no conflict of interest.

## Supplementary Figures

**Figure S1.Schematic diagram of the σ^B^ operon and the activation of σ^B^.**

A) Genetic organisation of the σ^B^ operon and *rsbZ* in *C. difficile* R20291. The σ^B^ operon and *rsbZ* locus are controlled by σ^A^. B) Activation of σ^B^-dependent genes. RsbZ is believed to sense stress and dephosphorylate RsbV. RsbW preferentially binds to RsbV, which releases σ^B^ from the control of RsbW and subsequently directs RNA polymerases to σ^B^ promoter sites.

**Figure S2.Δ*rsbW* does not display any fitness defect in a non-stressed state.**

Growth curves of *C. difficile* strains *C. difficile* WT, Δ*rsbW* and Δ*rsbW*::*rsbW* grown for 12 h in A) BHI-S, B) TY and C) DMEM media. D) Growth profiles were analysed with the GrowthCurver R package to calculate the generation time. N=3, error bars indicate SD.

**Figure S3.Overexpression of *rsbW* does not induce cellular toxicity.**

A) Overnight cultures of *C. difficile* WT, Δ*rsbW* and Δ*rsbW*::*rsbW* were normalised to OD_600_ 0.1 and spot diluted onto BHI-S agar with/without thiamphenicol/anhydrotetracycline. Viable bacteria were enumerated to quantify B) bacterial fitness and toxicity of anhydrotetracycline and C) plasmid-loss mediated survival. N=3 (3 technical replicates/experiment). Error bars indicate SD, no significance was denoted with n.s. and significance with \**p-*value < 0.05 as determined by Mann-Whitney U test

**Figure S4.Δ*rsbW* does not confer increased tolerance to oxidative tolerance in agar.**

Overnight cultures of *C. difficile* WT, Δ*rsbW* and Δ*rsbW*::*rsbW* were spiked onto 0.4% TY agar and grown aerobically for 24 h. A) Image of the zone of inhibition between the top of bacterial growth and the agar. B) The zone of inhibition was measured for each tube and the distribution was graphically represented. Data analysis was conducted with a Student’s t-test. No significance was denoted with n.s.

**Figure S5.No alteration in growth is observed when *C. difficile* strains subjected to pH 4, 6 and 7.**A) Growth profile of *C. difficile* WT, Δ*rsbW* and Δ*rsbW*::*rsbW* grown in BHI-S media calibrated to pH 4, 6 and 7. B) At each timepoint, cultures were grown on BHI-S agar plates to enumerate viable colonies.

**Figure S6.Bacterial stress tolerance to ROS (H_2_O_2_) and RNS (sodium nitroprusside).**

*C. difficile* WT, Δ*rsbW* and Δ*rsbW*::*rsbW* were grown on bacterial lawns were exposed to 1, 2, 4 and 9.8 M of H_2_O_2_ on 10 mm disks. The diameter of the zone of inhibition was measured. Viable bacteria were enumerated from *C. difficile* strains grown on TY agar plates supplemented with 0, 200, 500 and 1000 mM sodium nitroprusside. Differential survival was calculated with fold change between the bacterial grown with and without SNP. N=3 (3 technical replicates/experiment). Error bars indicate SD, no significance was denoted with n.s and significance with * *p-*value < 0.05 as determined by Student’s t-test and Mann-Whitney U test.

**Figure S7.RsbW alters biofilm thickness at 72 h.**

The voxel depth from the confocal microscopy images of each biofilm was calculated in Fiji (ImageJ). N=3 (3 technical replicates/experiment). No significance was denoted with n.s and significance with * = *p-*value < 0.05. as determined by Mann-Whitney U test.

**Figure S8.Transcriptomic analysis of Δ*rsbW* vs WT**

Biological replicates of the RNAseq data for *C. difficile* WT and Δ*rsbW* were analysed with A) Principal component analysis and B) Correlation coefficient analysis.

**Figure S9.Δ*rsbW* does not significantly alter bacterial ability to swim, swarm and produce toxin.**Bacterial motility was measured on soft agar plates over 24 h and 48 h, A) 0.3% agar for swimming and B) 0.4% agar for swarming. N=3 (3 technical replicates/experiment) C) Planktonic bacterial cultures were grown for 10 h, and toxin concentration were examined using ELISA for toxin A or B. N=3 (3 technical replicates/experiment). No significance was denoted with n.s. as determined by Mann-Whitney U test.

**Figure S10.Mass spectrometry reveals constitutive expression of σ^B^-controlled rubrerythrins in Δ*rsbW*.**A) Raw protein lysates obtained from *C. difficile* WT, Δ*rsbW* and Δ*rsbW*::*rsbW* grown to early exponential and stationary phase in BHI-S or TY medium. Samples were run through 12.5% polyacrylamide gel and stained with Coomassie blue. Excised bands were at ∼21 kDa (denoted with white arrows) were digested and analysed by mass spectrometry, revealing B) a distribution of normalised total protein spectra and C) differential abundance of rubreythrins in both BHI-S and TY.

## References

1. Leffler, D. A. & Lamont, J. T. Clostridium difficile infection. N. Engl. J. Med. 372, 1539–1548 (2015).

2. Czepiel, J. et al. Clostridium difficile infection: review. Eur. J. Clin. Microbiol. Infect. Dis. 38, 1211–1221 (2019).

3. Guh, A. Y. et al. Trends in U.S. Burden of Clostridioides difficile Infection and Outcomes. N. Engl. J. Med. 382, 1320–1330 (2020).

4. Wilcox, M. H. et al. Impact of recurrent Clostridium difficile infection: Hospitalization and patient quality of life. J. Antimicrob. Chemother. 72, 2647–2656 (2017).

5. Vardakas, K. Z. et al. Treatment failure and recurrence of Clostridium difficile infection following treatment with vancomycin or metronidazole: A systematic review of the evidence. Int. J. Antimicrob. Agents 40, 1–8 (2012).

6. Yearsley, K. A. et al. Proton pump inhibitor therapy is a risk factor for Clostridium difficile-associated diarrhoea. Aliment. Pharmacol. Ther. 24, 613–619 (2006).

7. Jump, R. L. P., Pultz, M. J. & Donskey, C. J. Vegetative Clostridium difficile survives in room air on moist surfaces and in gastric contents with reduced acidity: A potential mechanism to explain the association between proton pump inhibitors and C. difficile-associated diarrhea? Antimicrob. Agents Chemother. 51, 2883–2887 (2007).

8. Wetzel, D. & McBride, S. M. The Impact of pH on Clostridioides difficile Sporulation and Physiology. Appl. Environ. Microbiol. 86, 1–13 (2019).

9. Waligora, A. J., Barc, M. C., Bourlioux, P., Collignon, A. & Karjalainen, T. Clostridium difficile cell attachment is modified by environmental factors. Appl. Environ. Microbiol. 65, 4234–8 (1999).

10. Hecker, M., Pané-Farré, J. & Völker, U. SigB-dependent general stress response in Bacillus subtilis and related gram-positive bacteria. Annu. Rev. Microbiol. 61, 215–236 (2007).

11. Benson, A. K. & Haldenwang, W. G. Bacillus subtilis sigma B is regulated by a binding protein (RsbW) that blocks its association with core RNA polymerase. Proc. Natl. Acad. Sci. 90, 2330–2334 (1993).

12. Voelker, U., Dufour, A. & Haldenwang, W. G. The Bacillus subtilis rsbU gene product is necessary for RsbX-dependent regulation of σ(B). J. Bacteriol. 177, 114–122 (1995).

13. Vijay, K., Brody, M. S., Fredlund, E. & Price, C. W. A PP2C phosphatase containing a PAS domain is required to convey signals of energy stress to the σ(B) transcription factor of Bacillus subtilis. Mol. Microbiol. 35, 180–188 (2000).

14. Chen, C. C., Lewis, R. J., Harris, R., Yudkin, M. D. & Delumeau, O. A supramolecular complex in the environmental stress signalling pathway of Bacillus subtilis. Mol. Microbiol. 49, 1657–1669 (2003).

15. Yang, X., Kang, C. M., Brody, M. S. & Price, C. W. Opposing pairs of serine protein kinases and phosphatases transmit signals of environmental stress to activate a bacterial transcription factor. Genes Dev. 10, 2265–2275 (1996).

16. Kazmierczak, M. J., Wiedmann, M. & Boor, K. J. Alternative Sigma Factors and Their Roles in Bacterial Virulence. Microbiol. Mol. Biol. Rev. 69, 527–543 (2005).

17. Kint, N. et al. The alternative sigma factor σB plays a crucial role in adaptive strategies of Clostridium difficile during gut infection. Environ. Microbiol. 19, 1933–1958 (2017).

18. Kint, N. et al. The σB signalling activation pathway in the enteropathogen Clostridioides difficile. Environ. Microbiol. 21, 2852–2870 (2019).

19. Benson, A. K. & Haldenwang, W. G. Regulation of σ(B) levels and activity in Bacillus subtilis. J. Bacteriol. 175, 2347–2356 (1993).

20. Boylan, S. A., Rutherford, A., Thomas, S. M. & Price, C. W. Activation of Bacillus subtilis transcription factor σ(B) by a regulatory pathway responsive to stationary-phase signals. J. Bacteriol. 174, 3695–3706 (1992).

21. Delumeau, O., Lewis, R. J. & Yudkin, M. D. Protein-protein interactions that regulate the energy stress activation of σB in Bacillus subtilis. J. Bacteriol. 184, 5583–5589 (2002).

22. Dufour, A. & Haldenwang, W. G. Interactions between a Bacillus subtilis anti-σ factor (RsbW) and its antagonist (RsbV). J. Bacteriol. 176, 1813–1820 (1994).

23. Boekhoud, I. M., Michel, A.-M., Corver, J., Jahn, D. & Smits, W. K. Redefining the Clostridioides difficile σ B Regulon: σ B Activates Genes Involved in Detoxifying Radicals That Can Result from the Exposure to Antimicrobials and Hydrogen Peroxide. mSphere 5, (2020).

24. Doan, T. H. D., Yen-Nicolaÿ, S., Bernet-Camard, M. F., Martin-Verstraete, I. & Péchiné, S. Impact of subinhibitory concentrations of metronidazole on proteome of Clostridioides difficile strains with different levels of susceptibility. PLoS One 15, 1–23 (2020).

25. Benson, A. K. & Haldenwang, W. G. Characterization of a regulatory network that controls σ(B) expression in Bacillus subtilis. J. Bacteriol. 174, 749–757 (1992).

26. Wise, A. A. & Price, C. W. Four additional genes in the sigB operon of Bacillus subtilis that control activity of the general stress factor σ(B) in response to environmental signals. J. Bacteriol. 177, 123–133 (1995).

27. Senn, M. M. et al. Molecular analysis and organization of the σB operon in Staphylococcus aureus. J. Bacteriol. 187, 8006–8019 (2005).

28. O’Byrne, C. P. & Karatzas, K. A. G. Chapter 5 The Role of Sigma B (σB) in the Stress Adaptations of Listeria monocytogenes: Overlaps Between Stress Adaptation and Virulence. Advances in Applied Microbiology vol. 65 (Elsevier Masson SAS, 2008).

29. Kint, N. et al. How the Anaerobic Enteropathogen Clostridioides difficile Tolerates Low O 2 Tensions. MBio 11, 1–17 (2020).

30. Fimlaid, K. A. et al. Global Analysis of the Sporulation Pathway of Clostridium difficile. PLoS Genet. 9, (2013).

31. Zhu, D., Bullock, J., He, Y. & Sun, X. Cwp22, a novel peptidoglycan cross-linking enzyme, plays pleiotropic roles in Clostridioides difficile. Environ. Microbiol. 21, 3076–3090 (2019).

32. Dapa, T. et al. Multiple factors modulate biofilm formation by the anaerobic pathogen Clostridium difficile. J. Bacteriol. 195, 545–555 (2013).

33. Dawson, L. F. et al. Characterisation of Clostridium difficile Biofilm Formation, a Role for Spo0A. PLoS One 7, (2012).

34. Anonye, B. O. et al. Probing clostridium difficile infection in complex human gut cellular models. Front. Microbiol. 10, 1–15 (2019).

35. Nale, J. Y., Chutia, M., Carr, P., Hickenbotham, P. T. & Clokie, M. R. J. ‘Get in Early’; Biofilm and Wax Moth (Galleria mellonella) Models Reveal New Insights into the Therapeutic Potential of Clostridium difficile Bacteriophages. Front. Microbiol. 7, 1383 (2016).

36. Nale, J. Y., Chutia, M., Cheng, J. K. J. & Clokie, M. R. J. Refining the galleria mellonella model by using stress marker genes to assess clostridioides difficile infection and recuperation during phage therapy. Microorganisms 8, 1–15 (2020).

37. Permpoonpattana, P. et al. Functional characterization of Clostridium difficile spore coat proteins. J. Bacteriol. 195, 1492–1503 (2013).

38. Castro-Córdova, P. et al. Entry of spores into intestinal epithelial cells contributes to recurrence of Clostridioides difficile infection. Nat. Commun. 12, 1–18 (2021).

39. Girinathan, B. P., Ou, J., Dupuy, B. & Govind, R. Pleiotropic roles of Clostridium difficile sin locus. PLoS Pathogens vol. 14 (2018).

40. Ciftci, Y., Girinathan, B. P., Dhungel, B. A., Hasan, M. K. & Govind, R. Clostridioides difficile SinR’ regulates toxin, sporulation and motility through protein-protein interaction with SinR. Anaerobe 59, 1–7 (2019).

41. Dawson, L. F. et al. Extracellular DNA, cell surface proteins and c-di-GMP promote biofilm formation in Clostridioides difficile. Sci. Rep. 11, 1–21 (2021).

42. Purcell, E. B. et al. A Nutrient-Regulated Cyclic Diguanylate Phosphodiesterase Controls Clostridium difficile Biofilm and Toxin Production during Stationary Phase. Infect. Immun. 85, 1–17 (2017).

43. Purcell, E. B., McKee, R. W., McBride, S. M., Waters, C. M. & Tamayo, R. Cyclic diguanylate inversely regulates motility and aggregation in clostridium difficile. J. Bacteriol. 194, 3307–3316 (2012).

44. Bradshaw, W. J., Roberts, A. K., Shone, C. C. & Acharya, K. R. The structure of the S- layer of Clostridium difficile. J. Cell Commun. Signal. 12, 319–331 (2018).

45. Palma, M. et al. Salicylic acid activates sigma factor B by rsbU-dependent and - independent mechanisms. J. Bacteriol. 188, 5896–5903 (2006).

46. Inaoka, T., Matsumura, Y. & Tsuchido, T. SodA and manganese are essential for resistance to oxidative stress in growing and sporulating cells of Bacillus subtilis. J. Bacteriol. 181, 1939–1943 (1999).

47. Weiss, A., Lopez, C. A., Beavers, W. N., Rodriguez, J. & Skaar, E. P. Clostridioides difficile strain-dependent and strain-independent adaptations to a microaerobic environment. Microb. genomics 7, (2021).

48. Neumann-Schaal, M. et al. Tracking gene expression and oxidative damage of O 2 - stressed Clostridioides difficile by a multi-omics approach. Anaerobe 53, 94–107 (2018).

49. Reder, A., Gerth, U. & Hecker, M. Integration of σB activity into the decision-making process of sporulation initiation in Bacillus subtilis. J. Bacteriol. 194, 1065–1074 (2012).

50. Mccourt, J., O’Halloran, D. P., Mccarthy, H., O’Gara, J. P. & Geoghegan, J. A. Fibronectin-binding proteins are required for biofilm formation by community-associated methicillin-resistant Staphylococcus aureus strain LAC. FEMS Microbiol. Lett. 353, 157–164 (2014).

51. Mitchell, G. et al. SigB Is a Dominant Regulator of Virulence in Staphylococcus aureus Small-Colony Variants. PLoS One 8, 1–14 (2013).

52. Barketi-Klai, A., Hoys, S., Lambert-Bordes, S., Collignon, A. & Kansau, I. Role of fibronectin-binding protein A in Clostridium difficile intestinal colonization. Journal of Medical Microbiology vol. 60 1155–1161 at https://doi.org/10.1099/jmm.0.029553-0 (2011).

53. Dhungel, B. A. & Govind, R. Spo0A Suppresses sin Locus Expression in Clostridioides difficile. mSphere 5, 1–10 (2020).

54. Deakin, L. J. et al. The Clostridium difficile spo0A gene is a persistence and transmission factor. Infect. Immun. 80, 2704–2711 (2012).

55. Dineen, S. S., McBride, S. M. & Sonenshein, A. L. Integration of metabolism and virulence by Clostridium difficile CodY. J. Bacteriol. 192, 5350–5362 (2010).

56. Pettit, L. J. et al. Functional genomics reveals that Clostridium difficile Spo0A coordinates sporulation, virulence and metabolism. BMC Genomics 15, 1–15 (2014).

57. Mandic-Mulec, I., Doukhan, L. & Smith, I. The Bacillus subtilis SinR protein is a repressor of the key sporulation gene spo0A. J. Bacteriol. 177, 4619–4627 (1995).

58. Slater, R. T., Frost, L. R., Jossi, S. E., Millard, A. D. & Unnikrishnan, M. Clostridioides difficile LuxS mediates inter-bacterial interactions within biofilms. Sci. Rep. 9, 1–15 (2019).

59. Goh, S., Ong, P. F., Song, K. P., Rily, T. V. & Chang, B. J. The complete genome sequence of Clostridium difficile phage φC2 and comparisons to φCD119 and inducible prophages of CD630. Microbiology 153, 676–685 (2007).

60. Meessen-Pinard, M., Sekulovic, O. & Fortier, L. C. Evidence of in vivo prophage induction during clostridium difficile infection. Appl. Environ. Microbiol. 78, 7662–7670 (2012).

61. Schubert, R. A., Dodd, I. B., Egan, J. B. & Shearwin, K. E. Cro’s role in the CI-Cro bistable switch is critical for λ’s transition from lysogeny to lytic development. Genes Dev. 21, 2461–2472 (2007).

62. Groth, A. C. & Calos, M. P. Phage integrases: Biology and applications. J. Mol. Biol. 335, 667–678 (2004).

63. Tao, L., Wu, X. & Sun, B. Alternative sigma factor σH modulates prophage integration and excision in staphylococcus aureus. PLoS Pathog. 6, 1–11 (2010).

64. Hengge-Aronis, R. Signal Transduction and Regulatory Mechanisms Involved in Control of the σ S (RpoS) Subunit of RNA Polymerase. Microbiol. Mol. Biol. Rev. 66, 373–395 (2002).

65. Lavey, N. P., Shadid, T., Ballard, J. D. & Duerfeldt, A. S. Clostridium difficile ClpP Homologues are Capable of Uncoupled Activity and Exhibit Different Levels of Susceptibility to Acyldepsipeptide Modulation. ACS Infect. Dis. 5, 79–89 (2019).

66. Michel, A. et al. Global regulatory impact of ClpP protease of Staphylococcus aureus on regulons involved in virulence, oxidative stress response, autolysis, and DNA repair. J. Bacteriol. 188, 5783–5796 (2006).

67. Krüger, E., Witt, E., Ohlmeier, S., Hanschke, R. & Hecker, M. The Clp proteases of Bacillus subtilis are directly involved in degradation of misfolded proteins. J. Bacteriol. 182, 3259–3265 (2000).

68. Fagan, R. P. & Fairweather, N. F. Clostridium difficile has two parallel and essential sec secretion systems. J. Biol. Chem. 286, 27483–27493 (2011).

69. Rocha, E. R., Tzianabos, A. O. & Smith, C. J. Thioredoxin reductase is essential for thiol/disulfide redox control and oxidative stress survival of the anaerobe Bacteroides fragilis. J. Bacteriol. 189, 8015–8023 (2007).

70. Baban, S. T. et al. The Role of Flagella in Clostridium difficile Pathogenesis: Comparison between a Non-Epidemic and an Epidemic Strain. PLoS One 8, (2013).

71. Edwards, A. & McBride, S. Determination of the in vitro Sporulation Frequency of Clostridium difficile. Bio-Protocol 7, 1–8 (2017).

72. Langmead, B. & Salzberg, S. L. Fast gapped-read alignment with Bowtie 2. Nat. Methods 9, 357–359 (2012).

73. Quinlan, A. R. BEDTools: The Swiss-Army tool for genome feature analysis. Curr. Protoc. Bioinforma. 2014, 11.12.1-11.12.34 (2014).

74. Love, M. I., Huber, W. & Anders, S. Moderated estimation of fold change and dispersion for RNA-seq data with DESeq2. Genome Biol. 15, 1–21 (2014).

75. Ramarao, N., Nielsen-Leroux, C. & Lereclus, D. The insect Galleria mellonella as a powerful infection model to investigate bacterial pathogenesis. J. Vis. Exp. e4392 (2012) doi:10.3791/4392.

76. Goodman, J. K., Zampronio, C. G., Jones, A. M. E. & Hernandez-Fernaud, J. R. Updates of the In-Gel Digestion Method for Protein Analysis by Mass Spectrometry. Proteomics 18, 1800236 (2018).

