## Supplementary figures and images for "The regulatory role of anti-sigma factor, RsbW, in *Clostridioides difficile* stress response, persistence and infection"

### supplemental figures

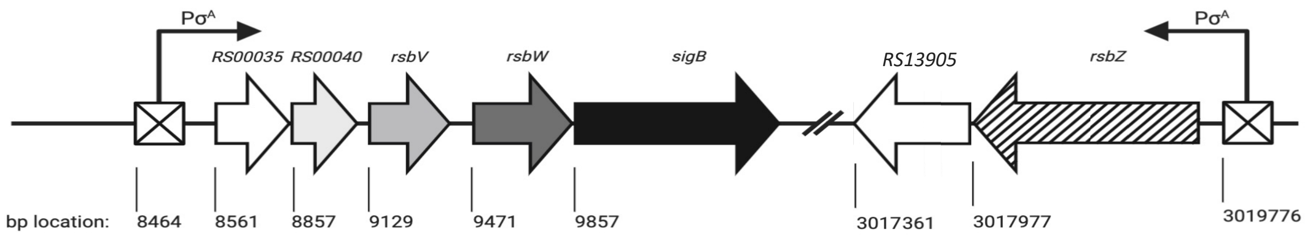

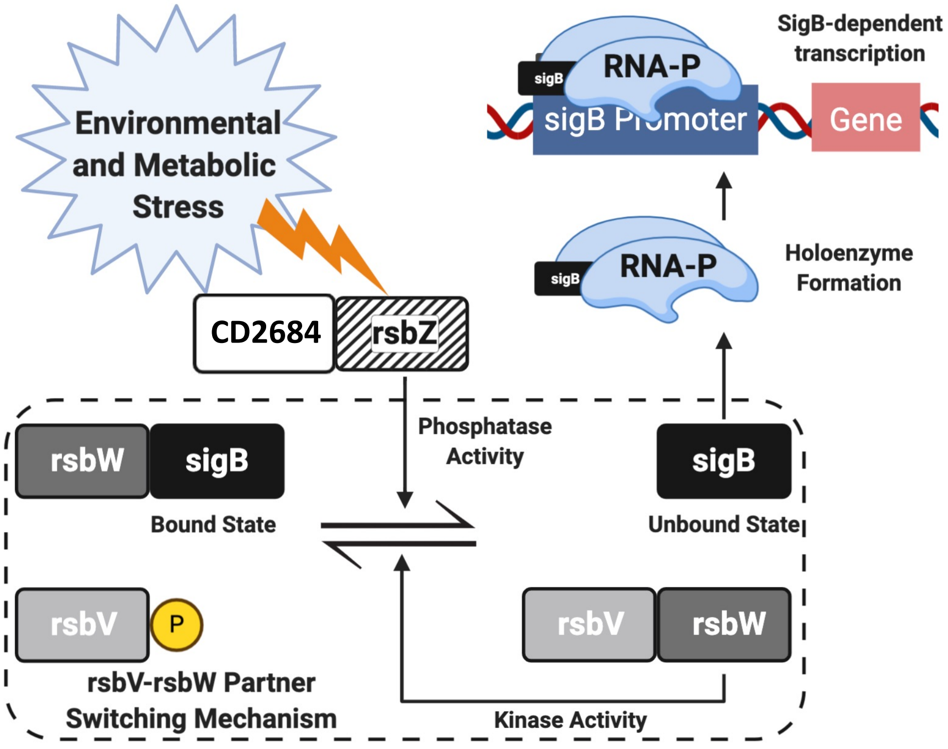

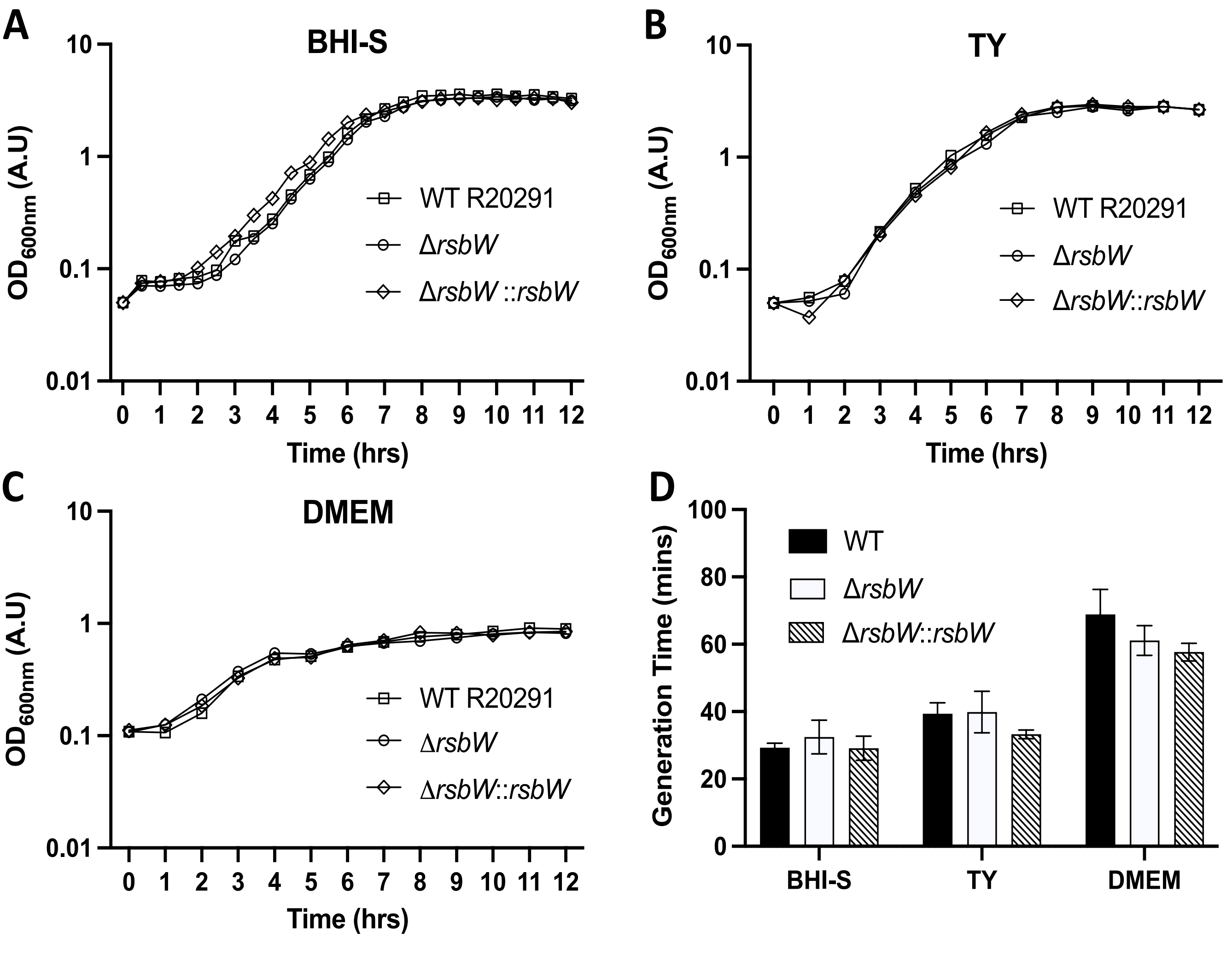

#### Figure S3

## A

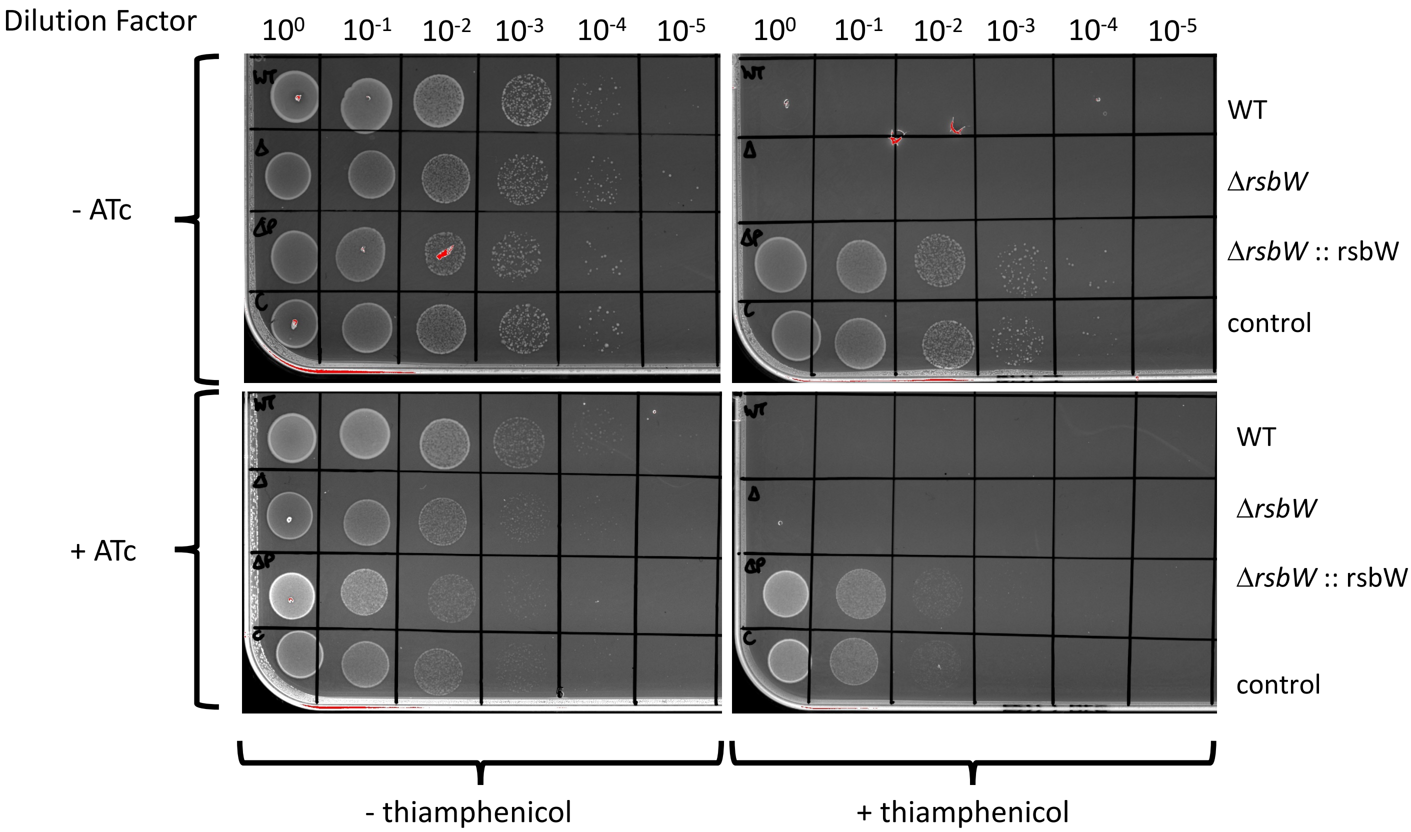


## B

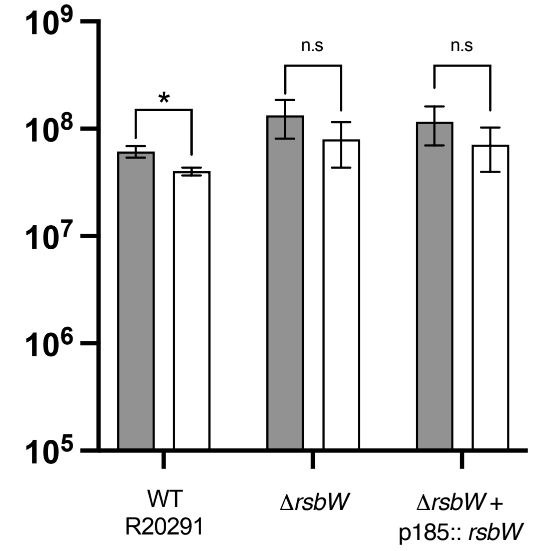


## C

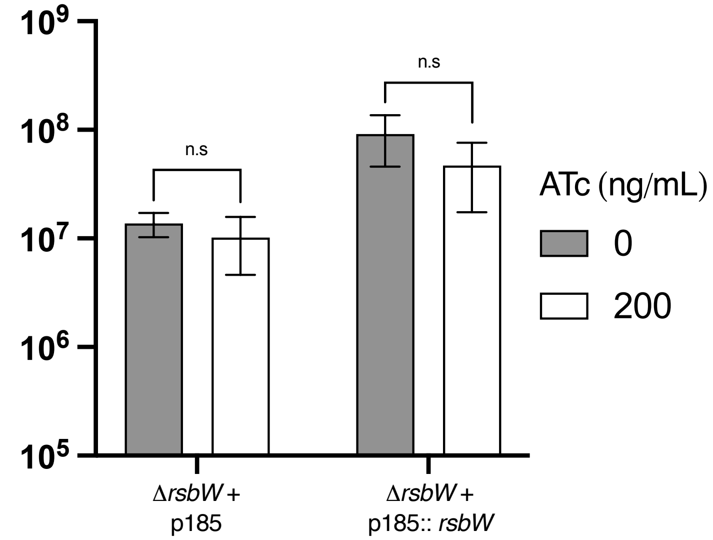

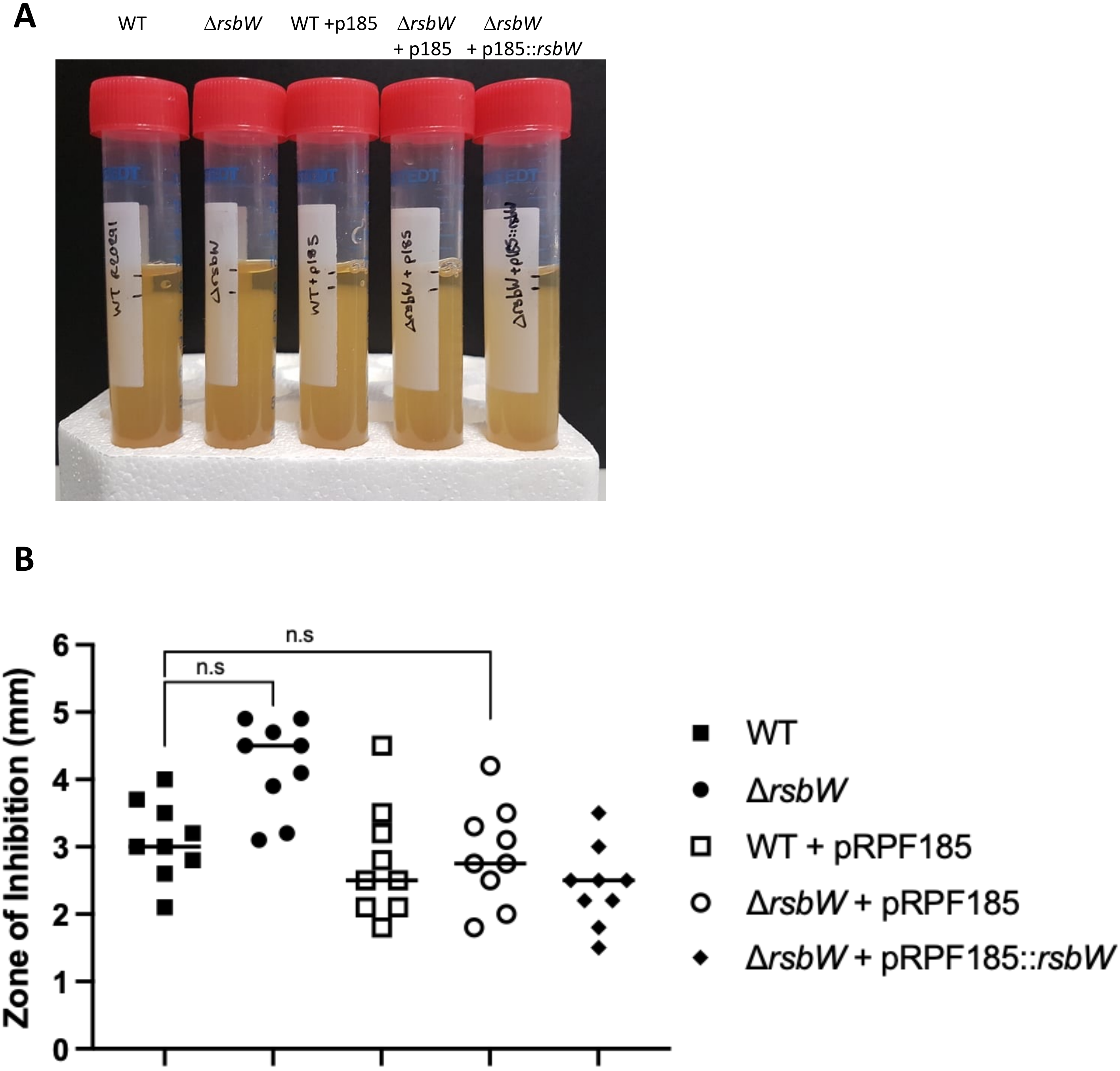

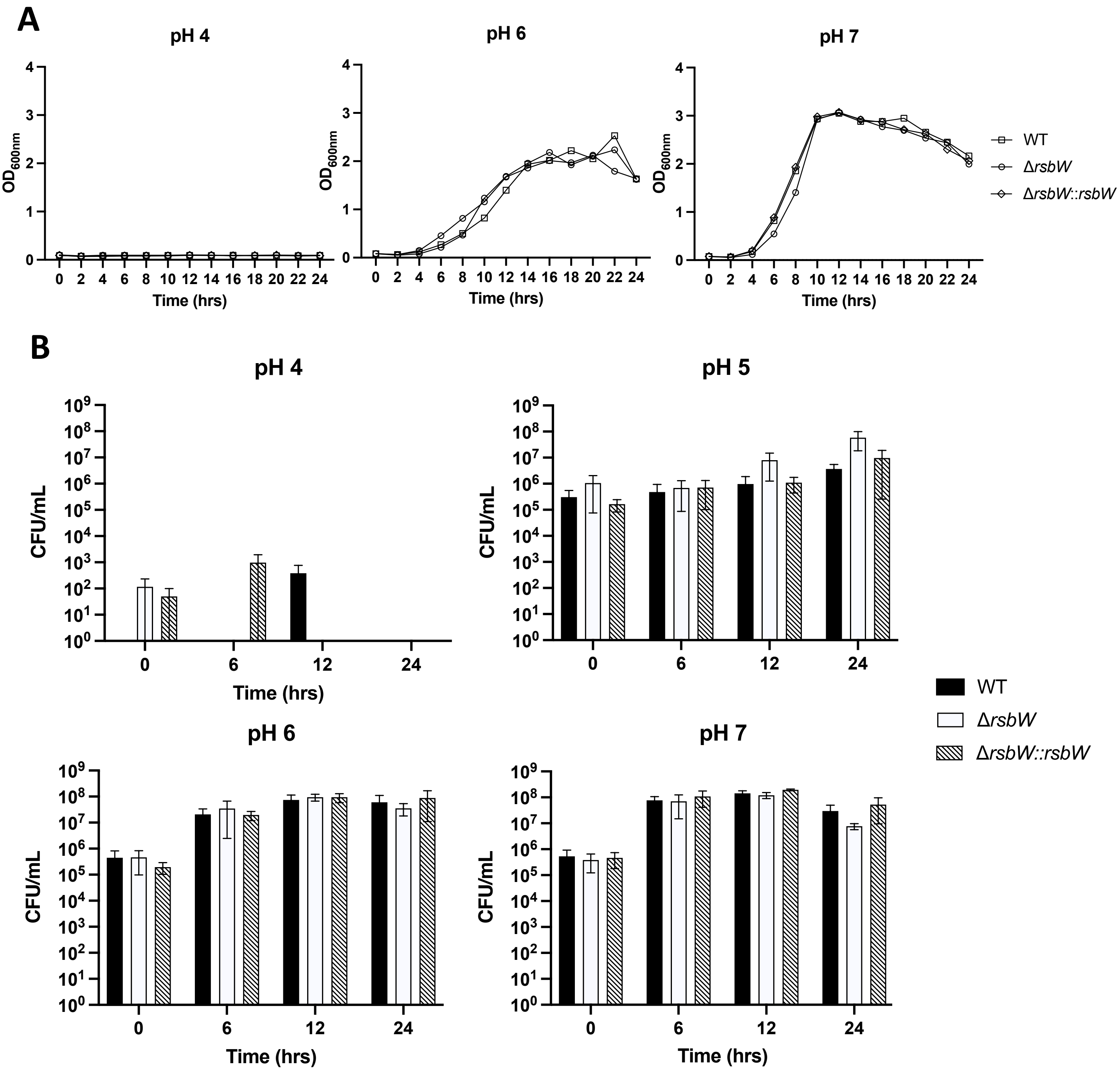

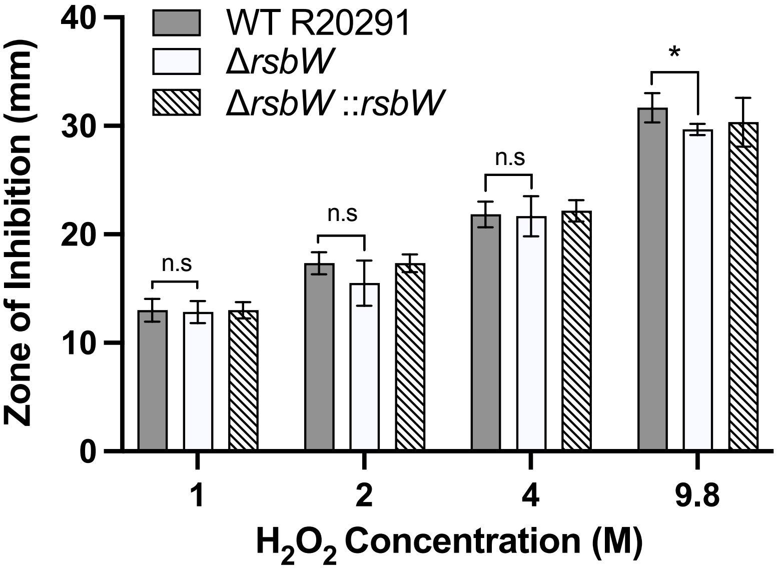

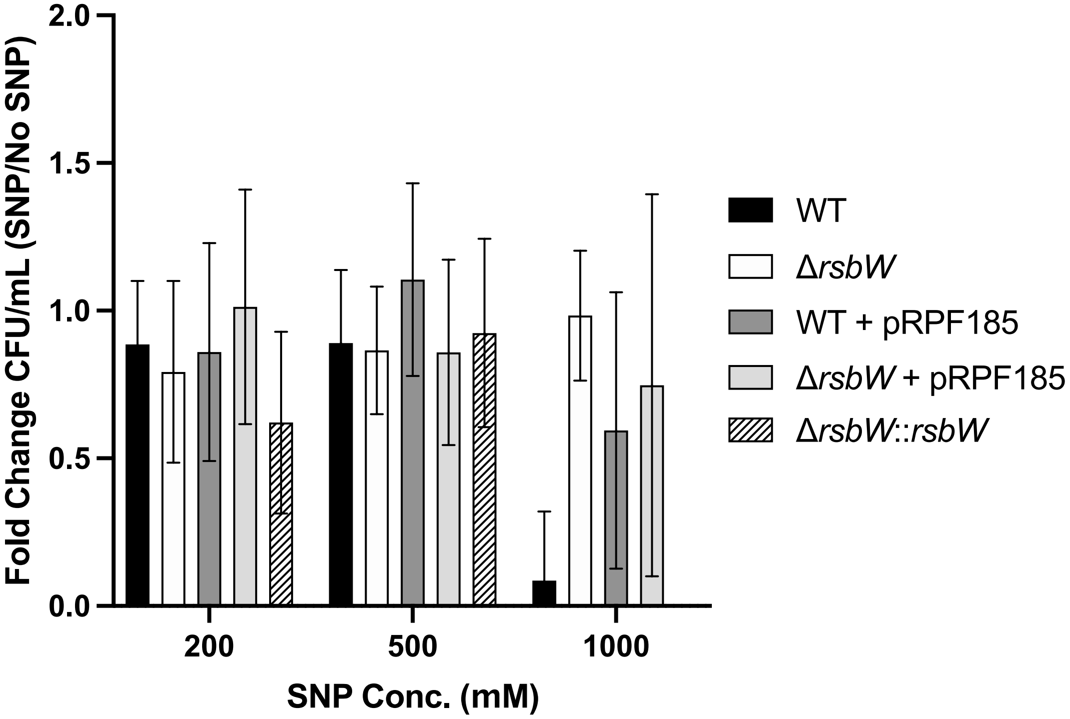

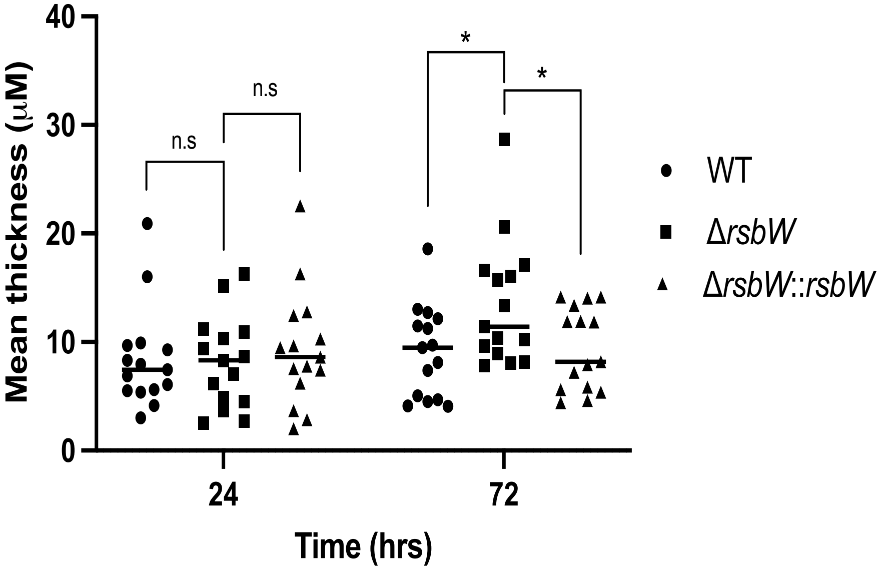

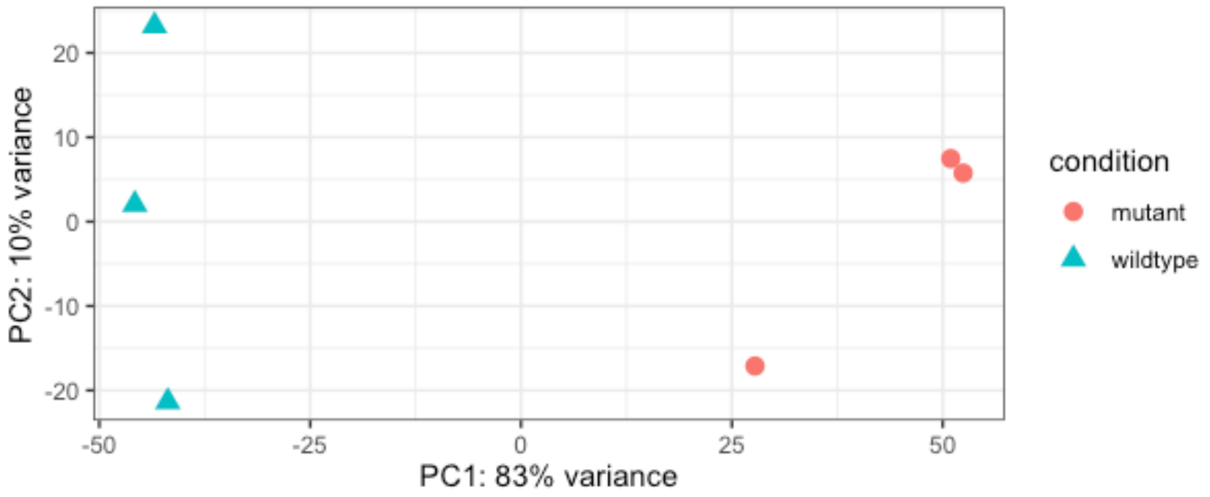

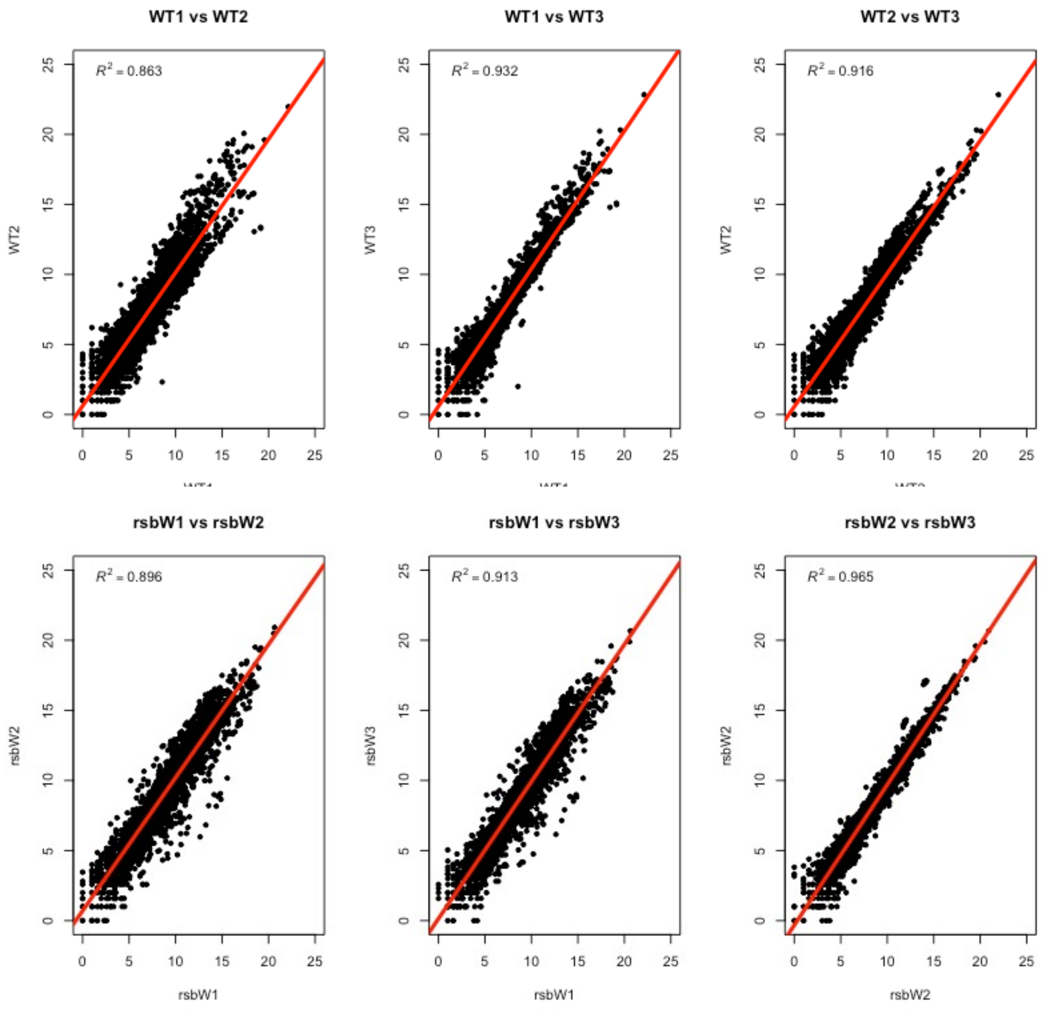

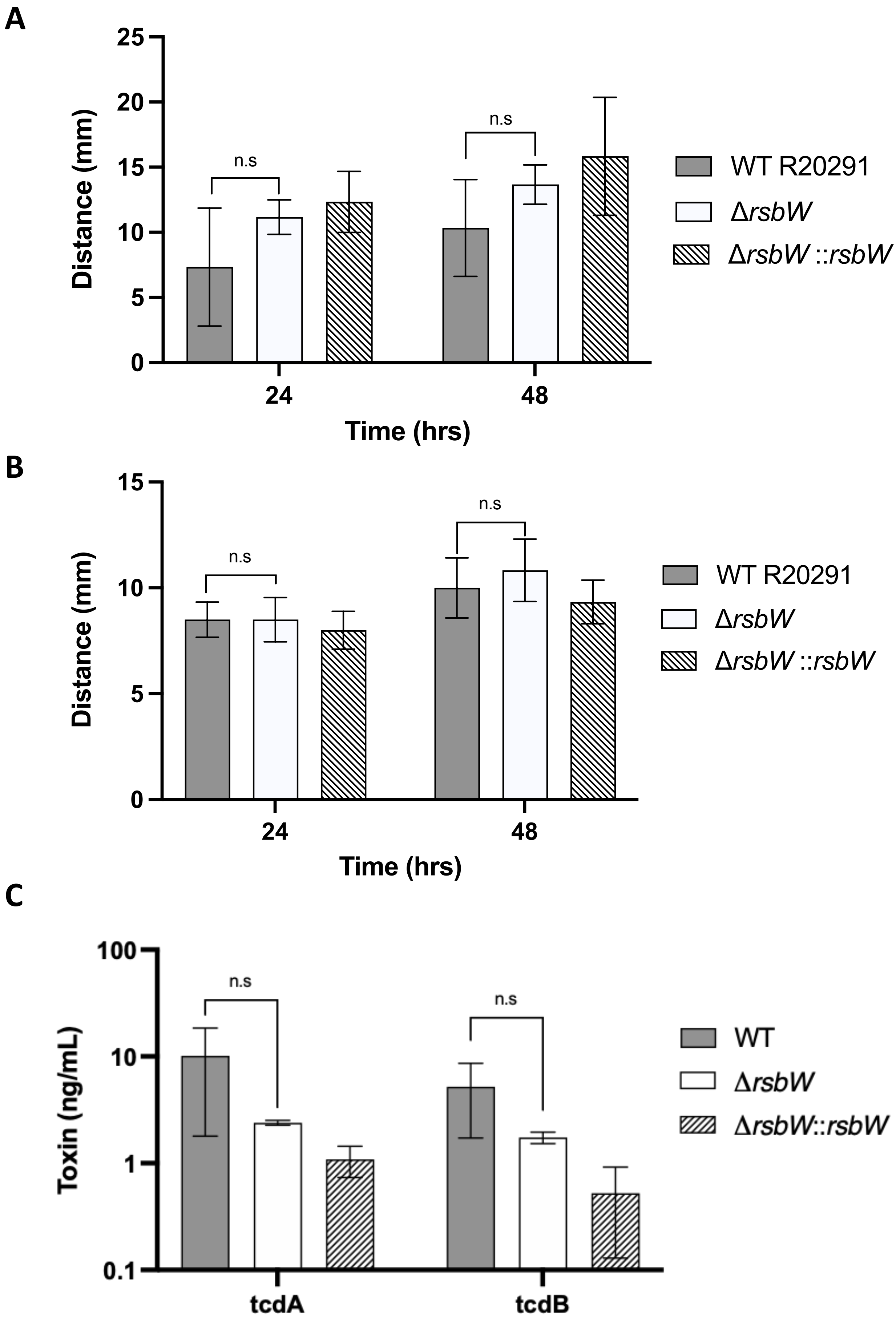

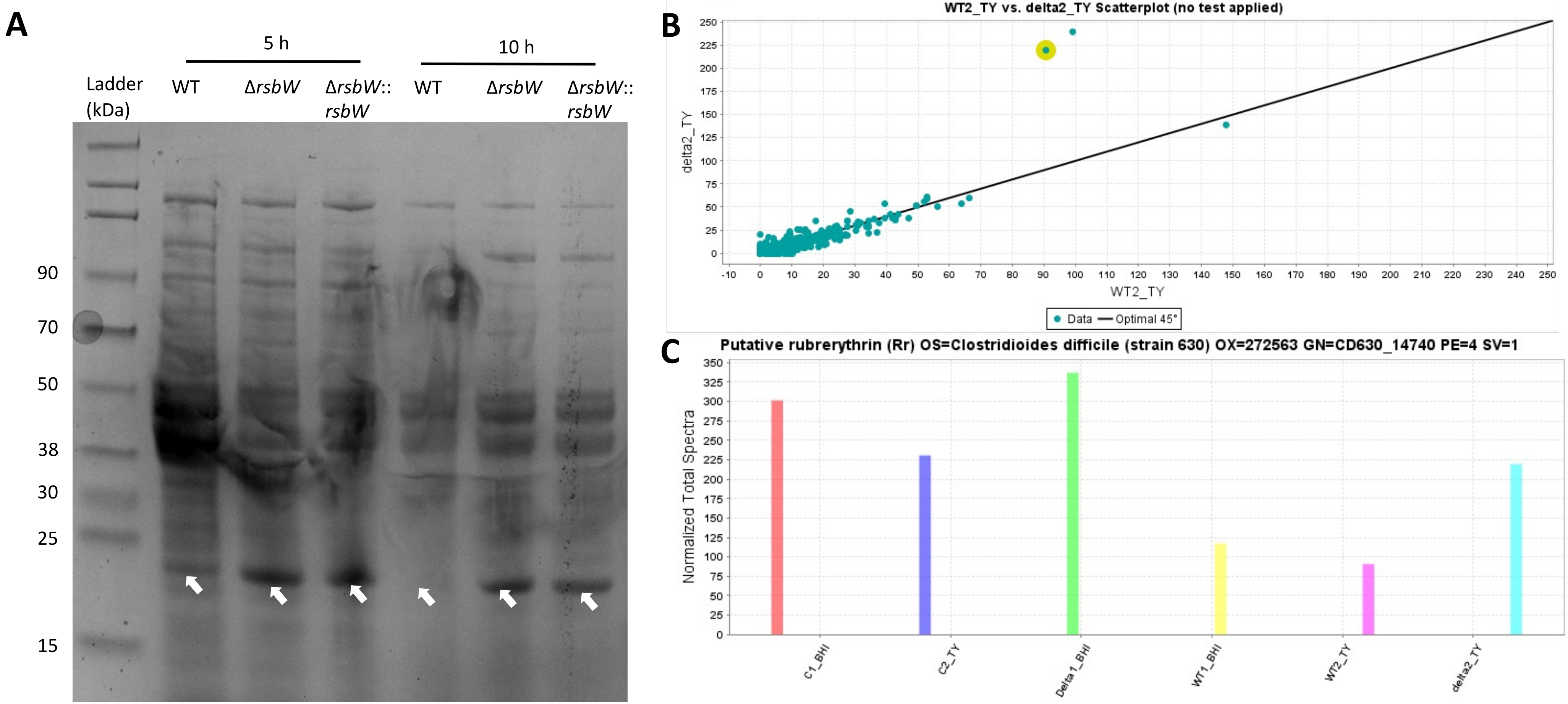
