## Supplementary material for "The regulatory role of anti-sigma factor, RsbW, in *Clostridioides difficile* stress response, persistence and infection"

**Supporting Material**

**Methods**

**Generation of Δ*rsbW***

*C. difficile* R20291Δ*rsbW* were generated using allelic exchange system (Cartman et al., 2012). Two allele exchange cassettes of approximately 500 bp were created from R20291 genomic DNA (isolated with DNeasy Blood and Tissue kit, Qiagen, Germany) flanking the *rsbW* gene. These homologous regions were amplified using primer couples F1_*rsbW*/R1_*rsbW* (upstream region) and F2_*rsbW*/F2_*rsbW* (downstream region) with Phusion High-Fidelity DNA polymerase (NEB, USA). Restriction sites *PmeI* and *EcoRI* were affixed at the END. PCR conditions were 95 °C for 5 mins, 35 cycles of: 95 °C for 30 secs, annealing temperature for 30 secs, extension for 72 °C for 1 min/kb and a final extension of 72 °C for 10 mins. One step ligation with Rapid DNA Ligation KIT (Roche, USA) was used to clone the two flanking regions into the PmeI site of pMTL-SC7315. This generated an in-frame deletion of the open reading frame of *rsbW* (nucleotides 10-401), named pMTL-SC7315-*rsbW*. The plasmid was conjugated into *E. coli* DH5α (Invitrogen, USA) by heatshock. Transformants were selected on LB agar supplemented with 25 µg/mL chloramphenicol, subcultured in LB broth supplemented with 12.5 µg/mL chloramphenicol. pMTL-SC7315-*rsbW* was purified and screened with PmeI restriction digestion (NEB, USA) and amplification with primer couples SC7-F1/SC7-R1. DNA sequence was confirmed with Sanger sequencing using SC7-F/SC7-R.

pMTL-SC7315-*rsbW* was conjugated into *E. coli* CA434 using electroporation and selected on LB agar supplemented with 25 µg/mL chloramphenicol. R20291 and CA434/pMTL-SC7315-*rsbW* were grown overnight in BHI-S and LB broth with 25 µg/mL chloramphenicol where appropriate. 1 mL of stationary culture of CA434/pMTL-SC7315-*rsbW* was pelleted and washed with 500 µL sterile PBS. The conjugal pellet was resuspended with 200 µL of R20291 stationary culture and plated onto pre-reduced BHI-S agar plates for incubated in anaerobic conditions for 24 h at 37 °C. Colonies were aggregated with 500 µL PBS and plated onto BHI-S supplemented with 250 µg/mL D-cycloserine, 8 µg/mL cefoxitin and 15 µg/mL thiamphenicol. Plates were incubated in anaerobic conditions for 24-72 hrs at 37 °C, transconjugants were restreaked onto similar plates and conditions. Single crossover events were purified, confirmed by PCR (RsbW_contr_F/SC7_R and RsbW_cont_R/SC7_R) and restreaked onto non-selective BHI-S agar in the same conditions for 96 hrs. Colonies were harvested in 500 µL sterile PBS, serially diluted to 10^-6^ and 100 µL spotted onto *C. difficile* Minimal Media supplemented with 50 µg/mL fluorocytosine. Second crossover mutants were isolated and patch plated onto BHI-S supplemented with D-cycloserine, cefoxitin and thiamphenicol. Fluorocytosine-resistant and thiamphenicol-sensitive clones were confirmed with PCR (F1­_*rsbW*/R2_*rsbW*) and Sanger sequencing.

**Supplementary Tables**

Table S1 Strains and plasmids used within this study

| **Strains** | **Function** | **Origin** |
| --- | --- | --- |
| *E. coli* |  |  |
| DH5α | Cloning Vector | Invitrogen, USA |
| CA434 | HB101(R702) Conjugation donor | (Purdy et al., 2002) |
| *C. difficile* |  |  |
| R20291 | Parental Strain | Lawley lab |
| Δ*rsbW* | R20291 with in-frame deletion of *rsbW* from | This study |
| R20291 + pRPF185 | WT Vector Control | This study |
| Δ*rsbW* + pRPF185 | Mutant Vector Control | This study |
| Δ*rsbW*::*rsbW* | Complementation of *rsbW* with pRPF185::*rsbW* | This study |
| Δ*spo0A* | Sporulation deficient mutant | Minton lab |
| **Plasmids** |  |  |
| pMTL-SC7315 | Allelic Exchange Vector | (Cartman et al., 2012) |
| pRPF185 | Inducible expression system. Tetracycline-inducible gusA. | (Fagan & Fairweather, 2011) |

Table S2

Oligonucleotides used within this study

| Primer | Sequence (5’ → 3’) | Function |
| --- | --- | --- |
| F1_*rsbW* | TTTTTTGTTTAAACTAAACAGCATAAATAAGGTTGT | Upstream flanking region with *PmeI* |
| R1_*rsbW* | GGGCTAGAATTCGTAATTTCCATCTTTATAGTCT | Upstream flanking region with *EcoRI* |
| F2_*rsbW* | GGGCTAGAATTCAATGACTAAATATTTAGGAGTTGA | Downstream flanking region with *EcoRI* |
| R2_*rsbW* | TTTTTTGTTTAAACTTGATATCCATAAGAAGCCTCC | Downstream flanking region with *PmeI* |
| SC7_F | GACGGATTTCACATTTGCCGTTTTGTAAACGAATTGCAGG | Sanger sequencing |
| SC7_R | AGATCCTTTGATCTTTTCTACGGGGTCTGACGCTCAGTGG | Sanger sequencing |
| Rsb_contr_F | TAGTAGGAAGCTCTGCTCTTATAGTAGC | Single crossover event |
| Rsb_contr_R | AATCTTCATACTCTATACTTCCAAAGTTACC | Single crossover event |
| F1_sigB_150bp | AGAGTCCCAAGAAGAATACAGGAA | RT-qPCR |
| R1_sigB_150bp | AGCCTCCATAGCCTCTAAAACA | RT-qPCR |
| F1_spo0A_146bp | TCAAAGCGCAATAAATCTAGGAGC | RT-qPCR |
| R1_spo0A_146bp | TCATTTGAGTCTCTTGAACTGGTCT | RT-qPCR |
| F1_sinR_153bp | AAAGGCAGGTTTACATCCAACAT | RT-qPCR |
| R1_sinR_153bp | GAGTTATCAACGCCTTCTGTTGT | RT-qPCR |
| F1_sigF_145bp | TGGAAGTAACTGTTGCCAGAGAA | RT-qPCR |
| R1_sigF_145bp | ACAACGCTCCTAACTAGACCT | RT-qPCR |
| F1_sigG_154bp | TGGGTCAAACAGAGATATTGGG | RT-qPCR |
| R1_sigG_154bp | AGCATATAAGGCACTACAAGTTAGA | RT-qPCR |
| F1_pil1A_153bp | GCTTTATCAGGCAGAGACTCCA | RT-qPCR |
| R1_pil1A_153bp | AGGCTAAGGTAGCAAGTGTTGA | RT-qPCR |
| F1_flgB_146bp | TGATGCTATGCCAAAAATAGAAGAA | RT-qPCR |
| R1_flgB_146bp | TCCATTTGCAAAACTTATCAAAGC | RT-qPCR |
| F1_slpA_151bp | AGCAAACTCAATAGTCGCAGC | RT-qPCR |
| R1_slpA_151bp | TGGAACTACTTATTCAACAGGTCT | RT-qPCR |
| F1_coll_bind_150bp | ACGACAAGTCCTACAATAGCTCC | RT-qPCR |
| R1_coll_bind _150bp | AGTGGTAAAGCCATCAGTGTCA | RT-qPCR |
| F1_ABC_tran _146bp | AGCAACAGGGTCACTCACAG | RT-qPCR |
| R1_ABC_tran_146bp | TGCCTACTACACAAAGTATTGCG | RT-qPCR |
| F1_vanR_155bp | GGGAAAGAAGTAGCATTAACACCG | RT-qPCR |
| R1_vanR_155bp | CGCCCTATATGAGCCATAACTGT | RT-qPCR |
| F1_glsA_149bp | ATGATGCTTCTGGGGAATTTGC | RT-qPCR |
| R1_glsA_149bp | AACTCCTGCGACGCTGTTAC | RT-qPCR |
| F1_norV_146bp | TCTGCACCAAATGCCATACAC | RT-qPCR |
| R1_norV_146bp | GGTTCATTTGGTTGGAGTGGTG | RT-qPCR |
| F1_nitro1_149bp | ATGAACGAATGGGGAAATGTTGC | RT-qPCR |
| R1_nitro1_149bp | ACAGCTAAAACTACAGTGCCTCC | RT-qPCR |
| F1_NADH_Per_152bp | GGTTATCCAGAAGTAGCTGAAGCA | RT-qPCR |
| R1_NADH_Per_152bp | TCAGTTGCACCATACTCAGCA | RT-qPCR |
| F1_FAD_OxiRed2_147bp | TGAAGGTGATAGTGCTCCTAGTG | RT-qPCR |
| R1_FAD_OxiRed2_147bp | GCATAAAAACCAGCTGCTCCA | RT-qPCR |

Table S3. Generation Time of *C. difficile* strains grown in pH 4-7

|  | **Mean generation time (h)+/- standard deviation (N=3)** | | |
| --- | --- | --- | --- |
|  | **WT** | **Δ*rsbW*** | **Δ*rsbW*::*rsbW*** |
| **pH 4** | No Growth | No Growth | No Growth |
| **pH 5** | 6.35 (± 0.85) | 4.38 (± 0.11) | 9.06 (± 1.76) |
| **pH 6** | 3.06 (± 0.57) | 3.31 (± 0.63) | 4.28 (± 1.77) |
| **pH 7** | 1.48 (± 0.38) | 1.44 (± 0.04) | 1.42 (± 0.57) |

**References**

Cartman, S. T., Kelly, M. L., Heeg, D., Heap, J. T., & Minton, N. P. (2012). Precise manipulation of the Clostridium difficile chromosome reveals a lack of association between the tcdC genotype and toxin production. *Applied and Environmental Microbiology*, *78*(13), 4683–4690. https://doi.org/10.1128/AEM.00249-12

Fagan, R. P., & Fairweather, N. F. (2011). Clostridium difficile has two parallel and essential sec secretion systems. *Journal of Biological Chemistry*, *286*(31), 27483–27493. https://doi.org/10.1074/jbc.M111.263889

Purdy, D., O’Keeffe, T. A. T., Elmore, M., Herbert, M., McLeod, A., Bokori-Brown, M., Ostrowski, A., & Minton, N. P. (2002). Conjugative transfer of clostridial shuttle vectors from Escherichia coli to Clostridium difficile through circumvention of the restriction barrier. *Molecular Microbiology*, *46*(2), 439–452. https://doi.org/10.1046/j.1365-2958.2002.03134.x
